# Single-cell transcriptomics uncover conserved molecular mechanisms and functional diversification in multilayered epithelia

**DOI:** 10.1101/2025.11.14.688286

**Authors:** Candice Merle, Mathilde Huyghe, Erica Kimber, Marisa M. Faraldo, Sophie Pantalacci, Marie Semon, Silvia Fre, Robin P. Journot

**Affiliations:** Institut Curie, PSL University, Sorbonne Université, CNRS UMR3215, Inserm U934, Genetics and Developmental Biology, 75005 Paris, France; Laboratory of Biology and Modeling of the Cell, Ecole Normale Supérieure de Lyon, CNRS, Inserm, UCBL, 69007, Lyon, France

**Keywords:** Epithelial stratification, p63-Notch axis, Single-cell transcriptomics, Comparative genomics, Gene regulatory networks

## Abstract

Multilayered epithelia, including skin, cervix, thymus, and prostate, arise from all three germ layers and perform diverse physiological functions, yet share a conserved layered architecture in which a supportive basal layer underlies differentiated suprabasal cells. To determine whether this common organization reflects a shared genetic program or tissue-specific regulation, we assembled a comprehensive single-cell atlas of 14 murine Multilayered epithelia.

We identified a conserved p63-Notch axis that universally governs basal cell identity maintenance and suprabasal commitment. The p63-centered transcriptional program is shared, while Notch signaling recruits context-specific transcriptional modules to generate functional diversity. The layered architecture of epithelia spatially and temporally decouples conserved basal from context-specific suprabasal networks, compartmentalizing functional innovation across multilayered epithelia.

Comparative genomics traces the p63-Notch axis to the root of vertebrates over 500 million years ago, showing that it was repeatedly co-opted through the incorporation of newly evolved genes in the suprabasal domain to generate novel epithelial functions. Our work establishes that evolution of multilayered epithelia operates through modular, compartmentalized diversification of a conserved scaffold, providing a unifying principle for how complex tissues achieve functional innovation while preserving structural integrity.

## Introduction

Multilayered epithelia are among the most adaptable tissues in vertebrates, forming protective barriers, secretory glands, and immunological interfaces across organs such as the skin, mammary gland, and thymus^1–3^. Arising from all three embryonic germ layers, ectoderm, endoderm, and mesoderm^4–11^, these tissues perform diverse physiological roles, yet they share a remarkably similar architecture: a basal compartment of progenitors supports suprabasal layers that differentiate into specialized lineages. This paradox of conserved architecture despite functional divergence raises a fundamental question: do multilayered epithelia share a universal molecular program, or does each organ deploy its own regulatory mechanisms to build a similar structure?

Work in the epidermis has described complementary regulatory roles for p63, a member of the p53 family that maintains basal progenitors^12–16^, and Notch signaling, which promotes suprabasal commitment^17–22^. Similar patterns of p63 expression in basal cells have been reported in other multilayered epithelia, including the trachea, bladder, mammary gland, and thymus^13,23–25^, although functional evidence outside the epidermis remains limited. Recent observations from our lab in glandular epithelia suggest that the p63–Notch axis may operate more broadly^26,27^, but whether this axis represents a universal regulatory principle of epithelial stratification remains unknown.

Multilayered epithelia first arose in vertebrates with epidermal stratification^28^, coinciding with the emergence of p63^29,30^. This raises the possibility that a conserved molecular framework underlies stratification, yet direct evidence across tissues is lacking.

Here we address this gap by constructing a single-cell transcriptomic atlas of 14 mouse multilayered epithelia spanning diverse germ-layer origins and functions. By integrating these data with lineage tracing, genetic perturbations, and gene regulatory network (GRN) inference, we reveal the existence of a universal spatial and temporal decoupling of two regulatory networks: a conserved basal program that maintains progenitor identity, and a modular suprabasal program that drives tissue-specific functional diversity.

Comparative genomics demonstrates that this regulatory compartmentalization is reflected in evolution, as suprabasal cells preferentially expressing newly evolved, lineage-specific genes. This indicates that the architectural layering partitions the deployment of evolutionary innovations. Our work thus defines a unifying framework for understanding multilayered epithelia organogenesis, revealing how architectural robustness can be balanced with functional diversity.

## Results

### A single-cell transcriptomic atlas of 14 mouse multilayered epithelia

To systematically map transcriptional diversity across multilayered epithelia, we compiled a comprehensive single-cell atlas of over 500,000 cells from 14 mouse organs, covering stratified and pseudo-stratified epithelia of diverse germ-layer origins, ages, sexes, and genetic backgrounds (Figure S1A-D). This atlas integrates 146 publicly available single-cell RNA sequencing (scRNA-seq) datasets from the epidermis^31–33^, cornea^34–36^, oral mucosa^37–39^, thymus^40–42^, mammary gland (MG)^43–46^, vagina^47,48^, cervix^48–50^, salivary gland (SG)^51,52^, bladder^53–56^, trachea^57–59^, lacrimal gland (LG)^60–62^, esophagus^63–65^, prostate^66–69^, and urethra^70,71^ (Figure 1A).

**Figure 1.**
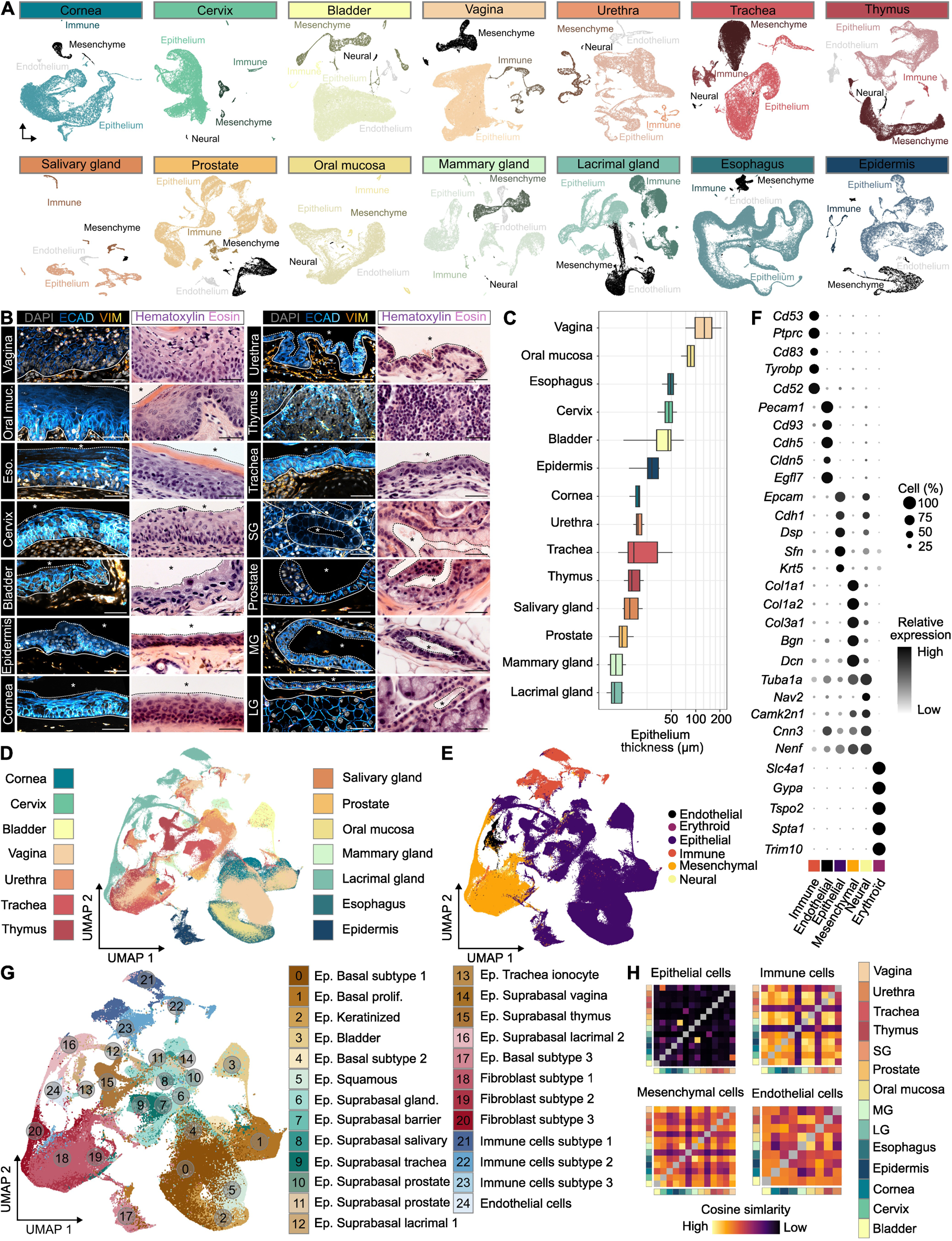
Integrated transcriptomic cell atlas of murine multilayered epithelia organs. ***A:*** *Uniform manifold approximation and projection (UMAP) of scRNA-seq data colored by organ and cell class. **B:** Confocal imaging of adult WT multilayered epithelia stained for Dapi, Ecadherin (ECAD, Cyan) and Vimentin (VIM, Orange) or Hematoxylin & Eosin staining. *: indicate lumen or external space; dotted line: indicate the interface between apical domain of epithelia and external space; continuous line: indicate the interface between epithelium and basement membrane. Scale bar: 50 µm. **C:** Quantification of epithelium thickness across multilayered epithelia (n=10 independent measurements). **D-E:** Integrated UMAP of scRNA-seq data colored by organ (D), and cell class (E). **F:** Dot plot showing cell class-level marker gene expression. **G:** UMAP of the multi-organ atlas, colored by major cell cluster, with shade and numbers representing cluster assignments. **H:** Cosine similarity between transcriptomes across organs, stratified by major cell class. To ensure robust representation, a threshold (≥100 cells) was applied for Immune, Mesenchymal, and Endothelial cell classes, retaining only tissues meeting this criterion. Rows and columns are color coded based on tissues*.

Among these multilayered epithelia, thickness varies extensively: protective epithelia such as the vagina, esophagus, and cornea form thick, multi-layered barriers, whereas glandular epithelia, including the mammary and salivary glands, are comparatively thin, typically comprising only two cell layers (Figure 1B, C). To correct for batch effects, background noise, and sex-related variation, we applied the Cluster Similarity Spectrum (CSS) integration method^72^ (Figure 1D, Figure S1E). The inclusion of three independent datasets per tissue allowed us to confirm that residual batch effects were minimal compared with genuine biological variation (Figure S1F–H). The resulting atlas includes not only epithelial cells but also stromal, immune, endothelial, neural, and erythroid populations (Figure 1E, F). Unsupervised clustering identified the major cell class across tissues, which were further resolved into 25 transcriptionally distinct transcriptomic clusters (Figure 1G; Figure S1I).

Remarkably, epithelial cells exhibited a much higher degree of transcriptional heterogeneity and organ specificity than stromal, endothelial, or immune cells (Figure 1E, G). To quantify this effect, we computed pairwise the cosine similarity between transcriptomes across all analyzed organs. Hierarchical clustering of these similarity scores confirmed that epithelial cells display the highest level of tissue specificity (Figure 1H).

Analyses of this atlas reveal that epithelial cells exhibit greater transcriptional heterogeneity and organ specificity than stromal or immune populations, establishing a high-resolution foundation to unravel the gene regulatory circuits that shape multilayered epithelia.

### Hierarchical basal–suprabasal transcriptional architecture is shared across organs

To resolve epithelial heterogeneity, we focused our analysis on the epithelial compartments of the integrated atlas. Tissue-specific sub clustering and marker-based annotation successfully recovered all known epithelial cell types across the 14 multilayered epithelia (Figure 2A; S2A). Remarkably, all tissues shared a basal cell (BC) population and one or more suprabasal populations of progenitors and differentiated cells, collectively referred to as suprabasal cells (SBCs).

**Figure 2.**
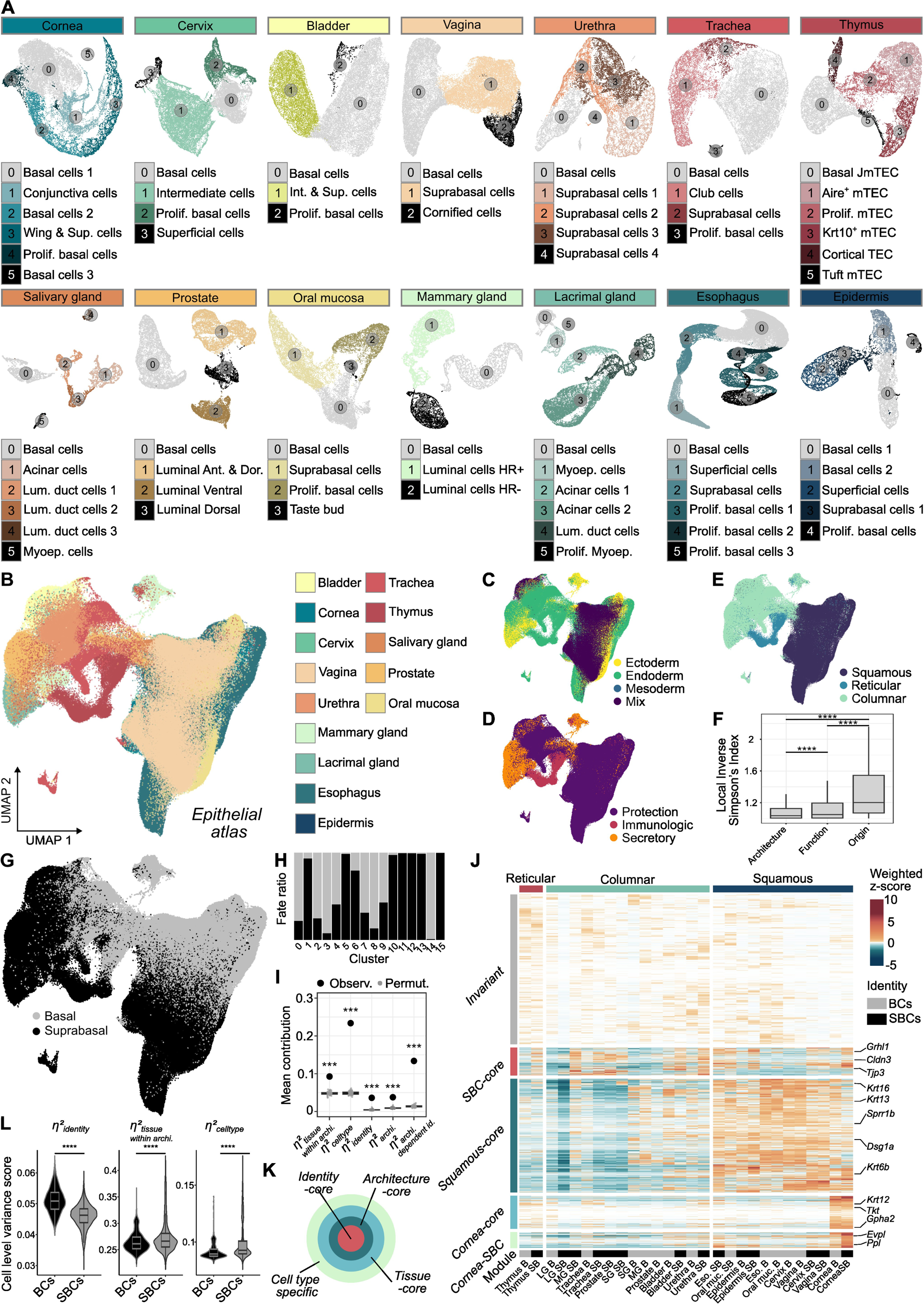
Identification of conserved spatial and transcriptomic organization in multilayered epithelia. ***A:*** *UMAP of epithelial cells scRNA-seq data colored by organ and cell cluster. Prolif.: Proliferative cells; Sup.: Suprabasal cells; mTEC: medullar Thymic epithelial cells; Lum.: Luminal; Myoep.: Myoepithelial; Ant.: Anterior; Dor.: Dorsal; HR: Hormone responsive. **B-E:** Integrated UMAP of epithelial scRNAseq data colored by organ (B), embryonic origin (C), physiological function (D) and tissue architecture (E). **F:** Local Inverse Simpson’s Index (LISI) scores for the epithelial integrated UMAP, reflecting the local diversity of cell classes in each clustering neighborhood, based on embryonic origin, physiological function or tissue architecture. Data displayed as boxplot. Non-parametric Wilcoxon test. **G:** Integrated UMAP colored by cell identity defined during analysis on individual organ datasets. **H:** Percentage of BCs and SBCs present in each cluster presented in Figure S2B. **I:** Boxplots show the null distributions of mean η² values obtained by permutation of Tissue and Identity labels (grey), while large black points indicate the observed mean η² for each variance component. Data displayed as boxplot, Permutation p-values (See Methods). **J:** Module decomposition of the cornea suprabasal transcriptome into Invariant, Suprabasal-core, Squamous-core, Cornea-core and Cornea-suprabasal modules (See Methods). **K:** Schematic Russian-doll decomposition of single cell transcriptome. **L:** Expression-weighted variance decomposition highlights divergent influences on basal and suprabasal cells. Expression-weighted η² values were aggregated at the single-cell level to quantify the relative influence of identity, function, tissue, and interaction terms on transcriptional variance. Data displayed as boxplot. Non parametric Wilcoxon test*.

After integration, epithelial cells failed to cluster by germ layer despite their diverse embryonic origins (Figure 2B, C). Instead, cells better grouped by physiological function and, most prominently, by tissue architecture (Figure 2D, E; Figure S2B). Classification into squamous, reticular, or columnar epithelia revealed the strongest separation in integrated space, quantitatively validated by Local Inverse Simpson’s Index (LISI) analysis (Figure 2F; see Methods). We next asked whether cells grouped by identity. In the integrated UMAP, BCs from all tissues coalesced into a continuous transcriptional domain, while SBCs formed discrete clusters defined by their architectural subtype: squamous versus reticular/columnar (Figure 2G, H; S2C-E).

To systematically dissect how cell identity, epithelial architecture, and tissue context shape these transcriptional differences, we decomposed per-gene variance into independent components using a linear contrast-based framework (Figure S2F, G; see Methods). This approach partitioned deviations from the global mean expression across tissues into biologically interpretable sources. It captured the distinct influence of cell identity (basal vs. suprabasal), epithelial architecture (squamous, columnar, reticular), and tissue-specific factors.

We next assigned genes to expression modules based on their dominant source of variance, ensuring that each module captured the main factor shaping its expression pattern (Figure S2H).

To validate that this hierarchical partitioning reflects genuine biological modularity rather than random structure, we performed permutation testing. Cell identity and tissue labels were randomly reassigned across samples 500 times (See Methods), and the proportion of variance (η²) explained by each factor was recalculated for each permutation. The observed η² values consistently surpassed those from the permuted datasets, confirming that the decomposition into *identity-*, *architecture-*, and *tissue-level* components captures statistically robust, non-random biological structure (Figure 2I). Supporting this, intra-module coherence analysis revealed strong correlation among genes within each module, further validating the decomposition’s robustness (Figure S2I).

The model yielded biologically interpretable insights. For instance, the transcriptional profile of corneal suprabasal cells could be reconstructed by integrating a *Suprabasal-core* module with *Squamous*, a *Cornea-core*, and *Cornea-specific suprabasal* modules (Figure 2J). This architecture suggests a Russian-doll-like organization of cell type transcriptomes: core identity modules form the innermost layer, with progressively specialized modules, linked to architecture, tissue, and cell type, forming concentric outer layers (Figure 2K).

To assess how transcriptomic variation differs between epithelial compartments, we applied the variance decomposition framework to individual cells grouped by identity. This analysis revealed a striking asymmetry. In basal cells, variance was predominantly explained by identity-related effects, suggesting that their transcriptional program is relatively stable across tissues. In contrast, suprabasal cells redistributed variance toward tissue- and cell type–specific components, indicating context-dependent diversification (Figure 2L).

Together, these analyses reveal a hierarchical organization of epithelial transcriptomes: a shared program maintains basal cell identity across all tissues, while suprabasal programs progressively diversify in an architecture- and tissue-specific manner.

### A shared p63-centered network defines basal identity and supports tissue-specific programs

The strong identity-driven transcriptional similarity of basal cells across tissues prompted us to ask whether a common GRN maintains basal identity in all multilayered epithelia. Analysis of the integrated basal compartments revealed a transcriptionally homogeneous population expressing canonical basal markers (*Krt5*, *Krt14*, *Krt15*) (Figure 3A, B), including a subset of proliferative cells (Figure S3A–B). The fraction of cycling BCs varied across organs, likely reflecting intrinsic differences in epithelial turnover (Figure S3B).

**Figure 3.**
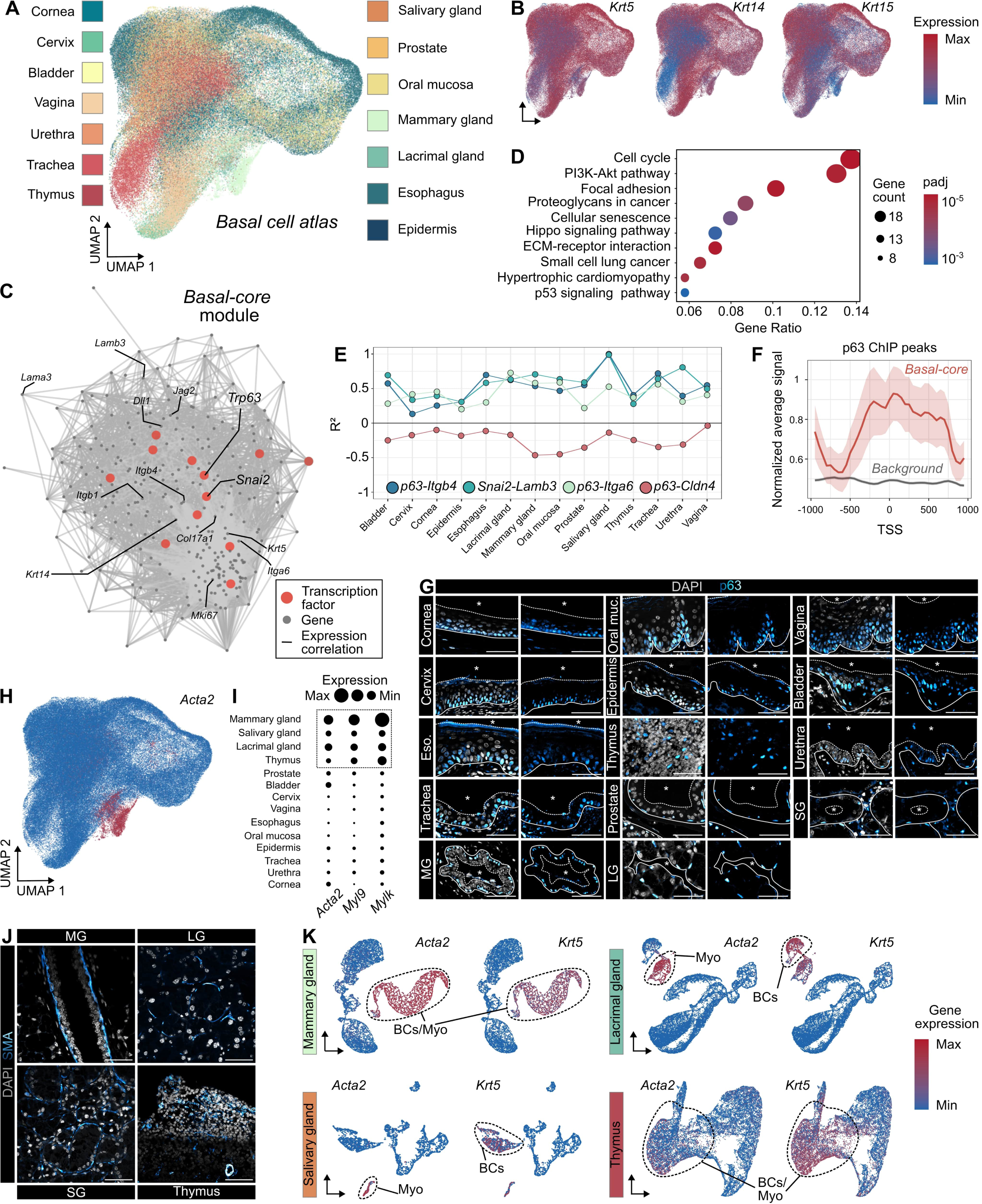
Characterization of the basal compartment: similarities and specialization across tissues. ***A:*** *Integrated UMAP of basal cells scRNAseq data colored by tissue. **B:** Integrated UMAP of scRNA-seq data colored by expression level of Krt5, Krt14 and Krt15 in individual cells. **C:** Basal-core GRN obtained by variance decomposition. Representation of the Basal-core as a correlation network of shared basal genes across tissues, highlighting transcription factors (red) as central hubs within the basal GRN. **D:** KEGG pathway enrichment across Basal-core module. **E:** Pairwise co-expression values for different selected correlating or anticorrelating gene pairs across the 14 tissues. **F**: Metagene profile of p63 ChIP-seq signal centered on transcription start sites (TSS) of basal-core genes (red) and background promoters (grey). The mean normalized signal (± standard error of the mean) is plotted in 50 bp bins across a ±1 kb window around the TSS. The p63 signal was capped at the 99.9th percentile and normalized for sequencing depth per chromosome. **G:** Confocal imaging of adult WT multilayered epithelia stained for Dapi and p63 (Cyan). *: indicate lumen or external space. **H:** Integrated UMAP of basal cells scRNAseq data colored by level of Acta2 gene expression. **I**: Dot plot showing percentage level of expression of genes associated with contractility (Acta2, Myl9 and Mylk) in the 14 murine multilayered epithelia. **J:** Confocal imaging of adult WT mouse section of Mammary, Lacrimal, Salivary gland and Thymus stained for Dapi (grey) and α-SMA (Cyan). **I:** UMAP of epithelial cells scRNA-seq data from Mammary, Salivary, Lacrimal gland and Thymus, colored by expression of Krt5 or Acta2 in individual cells. *: indicate lumen or external space; dotted line: indicate the interface between apical domain of epithelia and external space; continuous line: indicate the interface between epithelium and basement membrane. In G and J, scale bar: 50 µm*.

Using our previously described variance decomposition approach (Figure 2K), we identified ∼270 genes consistently enriched in basal cells, defining the *Basal-core* module (Figure 3C). Functional enrichment linked this module to cell-matrix adhesion, proliferation, and PI3K and Hippo pathways (Figure 3D), reflecting a coordinated regulatory program that underlies the defining structural and proliferative features of BCs.

Correlation analysis revealed that the *Basal-core* module forms a densely interconnected network of transcription factors and basal markers. *Trp63* and *Snai2* are at the core of the most consistently co-expressed network across tissues (Figure S3C). For example, *Snai2* correlated positively with *Dll1* and *Lamb3*, while *Trp63* correlated with the integrin genes *Itga6* and *Itgb4* (Figure 3E). Notably, p63-deficient mice fail to develop stratified epithelia^15,16^, whereas *Snai2*-deficient mice show only premature differentiation^73^, highlighting the functional hierarchy within this basal transcriptional backbone.

In addition, analysis of ΔNp63, the predominant *Trp63* isoform present in basal cells, using ChIP-seq data from mouse keratinocytes^74^ revealed a significant enrichment of p63 binding sites among *Basal-core* genes compared to the genomic background (Figure 3F; S3D). This included direct binding to *Krt5* and *Itgb4* and to ∼25% of the *Basal-core* genes overall (Figure S3E). Consistently, immunostaining for p63 and KRT5 across all 14 epithelia confirmed that the basal program is maintained at the protein level (Figure 3G; Figure S3F).

Having defined the conserved p63-centered Basal-core program, we asked whether basal cells exhibit tissue-specific variations. We further investigated whether these variations correspond to specialized functions. While some genes displayed tissue-restricted expression within basal compartments, these variations did not coalesce into coherent functional modules. Instead, a single, recurrent and coordinated program emerged; the formation of contractile basal myoepithelial cells.

These cells express smooth-muscle actin (*Acta2*, encoding for αSMA) alongside several contractility-associated genes such as *Myh11*, *Myl*9*, Mylk* and *Tpm2* (Figure 3H, I; S3G). SMA⁺ cells are present in the thymus, the mammary, lacrimal and salivary gland (Figure 3J). Analysis of individual datasets revealed distinct patterns. In the mammary gland and thymus, all BCs uniformly expressed *Acta2*. In contrast, salivary and lacrimal glands contained two basal subtypes: a ductal BC population expressing only *Krt5*, and an acinar BC cluster co-expressing *Krt5* and *Acta2* (Figure 3K).

These analyses define a shared, p63-centered GRN that establishes basal epithelial identity across multilayered epithelia. This core network forms a stable regulatory backbone upon which tissue-specific modules, such as the myoepithelial program, can be overlaid to generate functional diversity.

### Modular diversification drives organ-specific suprabasal programs and functions

While p63 governs a common basal program, the organization and tissue-specific variation of suprabasal transcriptional programs across organs remain poorly characterized. Integrative analysis of the suprabasal compartment revealed marked heterogeneity among SBC (Figure 4A), most apparent in keratins expression. Distinct combinations of simple (*Krt8/18/19*), mucosal (*Krt4/13/16*), or cornified (*Krt1/10/6a*) keratins contrast sharply with the uniform *Krt5/14* expression typical of basal cells (Figure 3B; 4B). Quantitative comparison of transcriptomic signatures revealed greater cross-tissue divergence among suprabasal than basal cells (Figure 4C), indicating that diversification is concentrated in this compartment.

**Figure 4.**
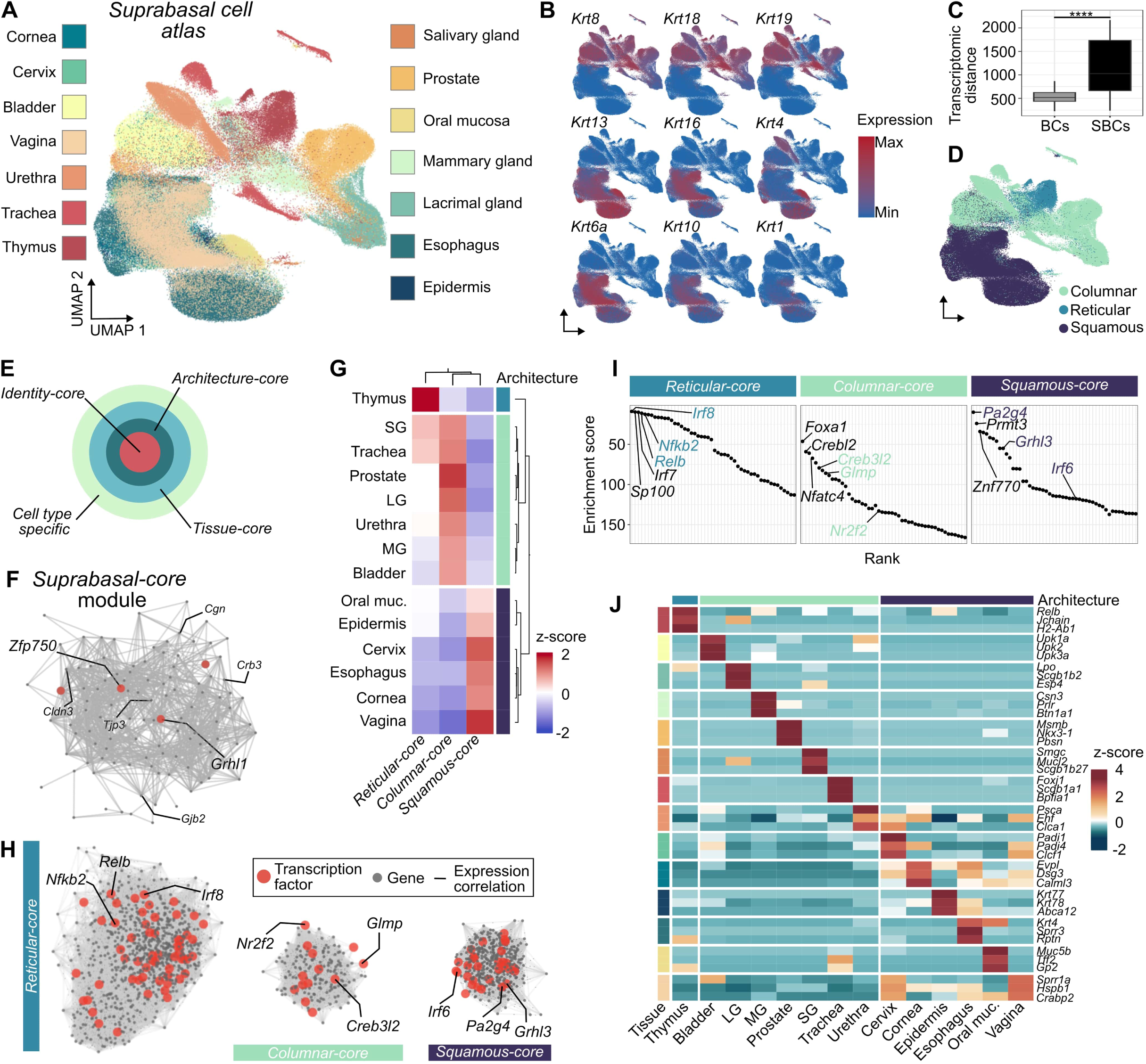
Diversification of the suprabasal transcriptome in relation to organ function. ***A:*** *Integrated UMAP of suprabasal cells scRNAseq data colored by tissue. **B:** Integrated UMAP of scRNA-seq data colored by expression level of keratins in individual cells. **C:** Euclidian distance across Basal and Suprabasal pseudo bulk samples. Data displayed as boxplot. Non parametric Wilcoxon test. **D:** Integrated UMAP of suprabasal cells scRNAseq data colored by tissue architecture (Columnar, Squamous, Reticular). **E:** Schematic Russian-doll decomposition of single cell transcriptome of suprabasal cells. **F:** Correlation network of the Suprabasal core module, highlighting transcription factors (red) as central hubs within the suprabasal GRN. **G:** Sparse regression–derived module weights are shown as a z-scored heatmap. Rows correspond to tissues, grouped by architectural class (Squamous, Columnar, Reticular), and columns correspond to architectural core modules. Color scale indicates relative module activity. **H:** Correlation network of the architecture core modules, highlighting transcription factors (red) as central hubs within the suprabasal GRNs of the indicated tissues. **I**: Transcription factor motif enrichment across architectural modules. Enrichment analysis of transcription factor–binding motifs performed with the ChEA database on genes defining the Columnar, Reticular, and Squamous core modules. Each plot shows the ranked enrichment scores of transcription factors, with representative top regulators annotated. Colored annotated TFs correspond to genes that belongs to the module. **J:** Z-score expression of Tissue and Suprabasal cell specific genes (3 genes per module)*.

In the integrated space, cells segregated according to epithelial architecture; squamous, columnar, or reticular, as observed in the whole epithelial atlas (Figure 4D). To formalize this organization, we applied our variance decomposition model, which partitions gene expression into nested components of shared, architectural, and tissue-specific variance (Figure 4E). This analysis revealed a compact *Suprabasal-core* module, approximately 2.5-fold smaller than the *Basal-core*, enriched for adhesion and cytoskeletal genes regulated by Grainyhead-family factors (*Grhl1*) and *Zfp750* (Figure 4F, G; S4A, B).

On top of this shared core, architecture-specific modules encode epithelial type, and include key transcription factors previously implicated in these epithelia: *Irf6*^75^ and *Grhl3*^76^ in squamous epithelia; *Relb*^77^, *Irf8*^78^, and *Nfkb2*^79^ in reticular epithelium; and *Nr2f2*^80^ and *Creb3l2*^81^ in columnar epithelia (Figure 4G, H). The hierarchical structure inferred from variance decomposition was independently corroborated by unbiased motif enrichment analysis, which identified the same transcriptional regulators among the top enriched motifs within each module (Figure 4I, S4C).

Yet tissues with a given architecture retained distinctive signatures. In squamous epithelia, SBCs spanned a spectrum from flexible mucosal keratins (*Krt4/13*), supporting adaptable suprabasal layers, to rigid keratins (*Krt1/10)* and cornified envelope proteins *(Ivl, Lor*), forming robust protective barriers (Figure S4D, E). *Tissue-core* modules varied even more, reflecting local regulatory inputs that overlay organ- and cell type–specific effectors on top of shared architectural modules, such as *Nkx2-1* in tracheal SBCs or *Dll4* in the thymic epithelium (Figure S4F). At the periphery of the hierarchy, *Cell type-specific* modules implement organ functions: *caseins* in the mammary gland, *uroplakins* in the bladder, and *mucins* or *secretoglobins* in the salivary and lacrimal glands (Figure 4J).

Within the framework of our variance decomposition, suprabasal transcriptomes mirror the hierarchical “Russian-doll” organization of basal cells. However, they exhibit markedly greater diversification, layering architectural and tissue-specific programs that encode organ-specific epithelial functions. Together with the conserved basal program, this modular organization allows multilayered epithelia to balance structural cohesion with organ-specific functional specialization.

### Notch signaling governs epithelial cell fate commitment across multilayered tissues

Having delineated the modular architecture of basal and suprabasal transcriptomes, we next sought to identify the molecular circuitry that connects them. Within the *Basal-core* GRN, we identified some Notch ligands (*Jag2*, *Dll1* and to a lesser extent *Jag1*) consistently co-expressed with *p63* across all multilayered epithelia (Figure 3H). By contrast, the direct Notch target and downstream effector *Hes1* was restricted to cells at the basal–suprabasal interface (Figure 5A). This sender-cell/receiver-cell topology implied that Notch may act as the molecular switch dictating fate commitment, consistent with its classical role as a universal arbiter of cell-fate decisions^82^.

**Figure 5.**
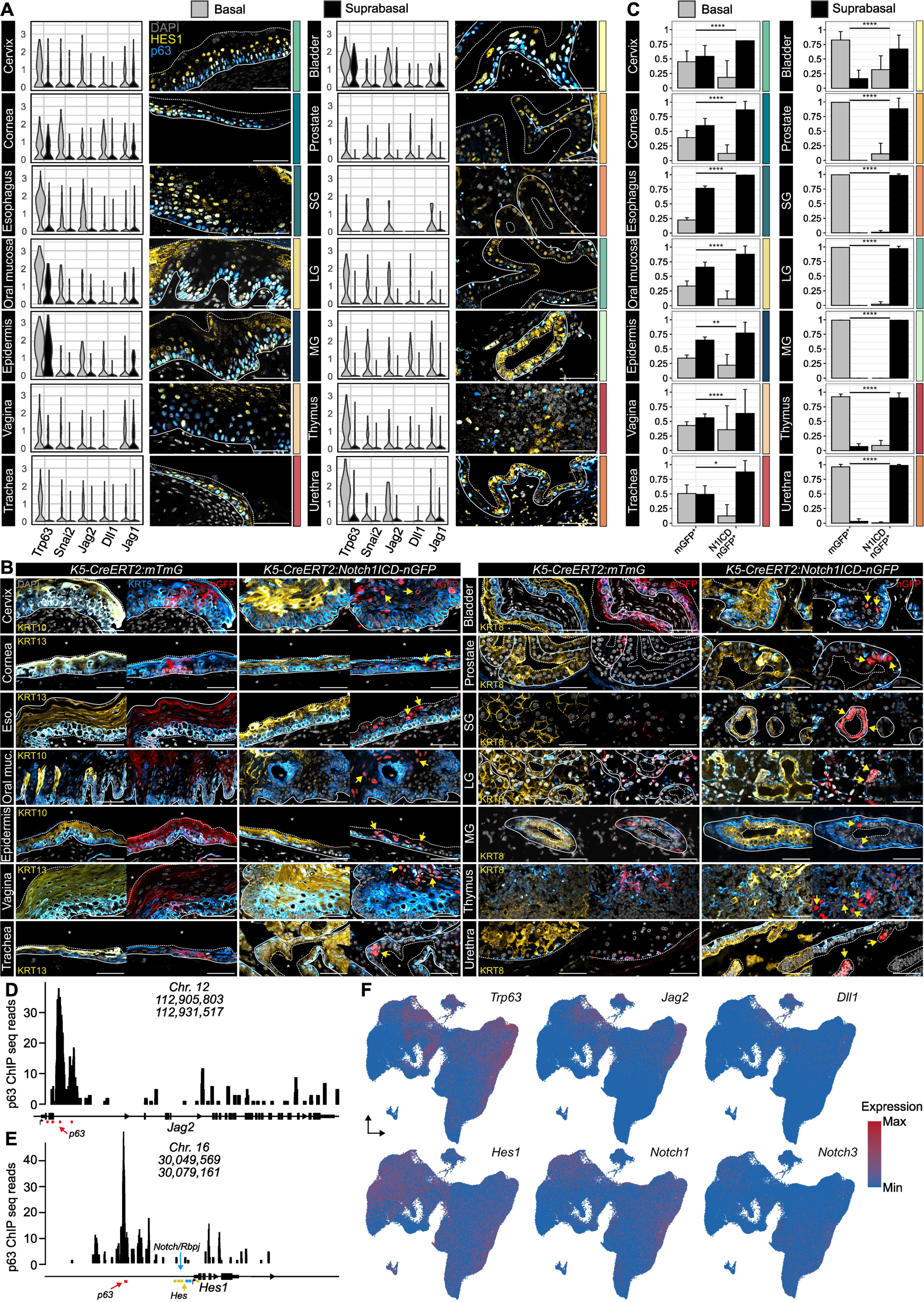
Notch signaling as a driver of suprabasal lineage commitment (Preliminary data) ***A:*** *Gene expression of Trp63, Snai2 and Notch ligands Dll1, Jag1 and Jag2 across BCs and SBCs of all 14 epithelia and confocal imaging of adult WT multilayered epithelia stained for Dapi, p63 and HES1. **B:** Confocal imaging of immunostaining for K5, SBCs marker and membrane-bound GFP (mGFP) in K5-Cre^ERT2^:mTmG (Control)or nuclear GFP (nGFP) in K5-Cre^ERT2^:N1ICD-nGFP (Notch GOF) epithelia after lineage tracing. **C:** Quantification of fate of mGFP^+^ or nGFP^+^ cells after 7 weeks of lineage tracing for mGFP^+^,1 week (Cervix, Cornea, Esophagus, Oral mucosa, Epidermis, Vagina and Trachea) or 7 weeks for nGFP^+^ (Bladder, Prostate, Salivary gland (SG), Lacrimal gland (LG), Mammary gland (MG), Thymus and Urethra). Data are displayed as Mean + s.d. Chi-squared test (n=3 biological replicates). **D-E:** ΔNp63 ChIP-seq read coverage across Jag2 (D) and Hes1 (E) loci (mm10 assembly). Signal tracks represent aligned ChIP-seq raw coverage for ΔNp63 in mouse keratinocyte. The predicted p63-binding site are colored in red. The approximate positions of two Notch/RBP-Jκ sites (blue)*^1^ *and four Hes-binding sites (yellow)*^2^ *are also shown. **F:** Integrated UMAP of scRNA-seq data colored by expression level of Trp63, Jag2, Dll1, Hes1, Notch1 and Notch3 in individual cells. *: indicate lumen or external space ; dotted line: indicate the interface between apical domain of epithelia and external space; continuous line: indicate the interface between epithelium and basement membrane. Scale bar: 50 µm*.

To test this model, we genetically forced the expression of constitutive form of the *Notch1* receptor in BCs using *K5-CreERT2:N1ICD-nGFP* mice^83^, and compared them to *K5-CreERT2:mTmG* controls^84^. Across all 14 tissues examined, Notch activation induced a basal-to-suprabasal conversion. Genetically traced mutant cells exited the basal layer and acquiring suprabasal or luminal markers, whereas control cells predominantly retained a basal cell identity (Figure 5B, C; S5A, B). The stability of lineage converted cells depended on tissue context: in rapidly renewing epithelia (Figure S3B), SBCs, including *N1ICD*⁺, were shed following upward migration. By contrast, converted luminal cells persisted or expanded in glandular epithelia (Figure S5C–E). These experiments indicate that Notch is sufficient to trigger suprabasal or luminal cell commitment, although the outcome of the mutant cells depends on the local tissue context and the turnover of SBCs in each tissue.

We next investigated how Notch is wired into the basal network. ΔNp63 ChIP-seq showed p63 occupancy at the *Jag2* promoter (overlapping previously mapped p63 motifs^85–87^) and at the *Hes1* promoter (Figure 5D–E), consistent with previous observations of p63 activating Notch ligand expression in BCs while repressing the downstream effector *Hes1*^19,87^. This dual regulation delineates two transcriptionally distinct compartments: *p63*⁺*Jag2*⁺ basal “signal-senders” and *p63*⁻*Hes1*⁺ suprabasal “signa-receivers” (Figure 5F). These results define a conserved p63–Notch module that links basal identity to suprabasal fate commitment.

Although previously recognized in the epidermis^17,22^, we now show that Notch signaling plays a decisive role in fate commitment across multiple homeostatic multilayered epithelia, including the vagina, cervix, urethra, oral mucosa, esophagus, and bladder. Thus, Notch emerges as a universal scaffold that coordinate progenitor maintenance and lineage segregation across multilayered epithelia.

### A conserved GRN underlies vertebrate epithelial stratification

Having established that murine multilayered epithelia share a common set of genes for stratification, rather than evolving tissue-specific solutions, we asked whether this core GRN reflects an evolutionarily conserved program. To address this, we integrated tissue morphology, single-cell transcriptomics, and gene conservation data across metazoans (Figure 6A).

**Figure 6.**
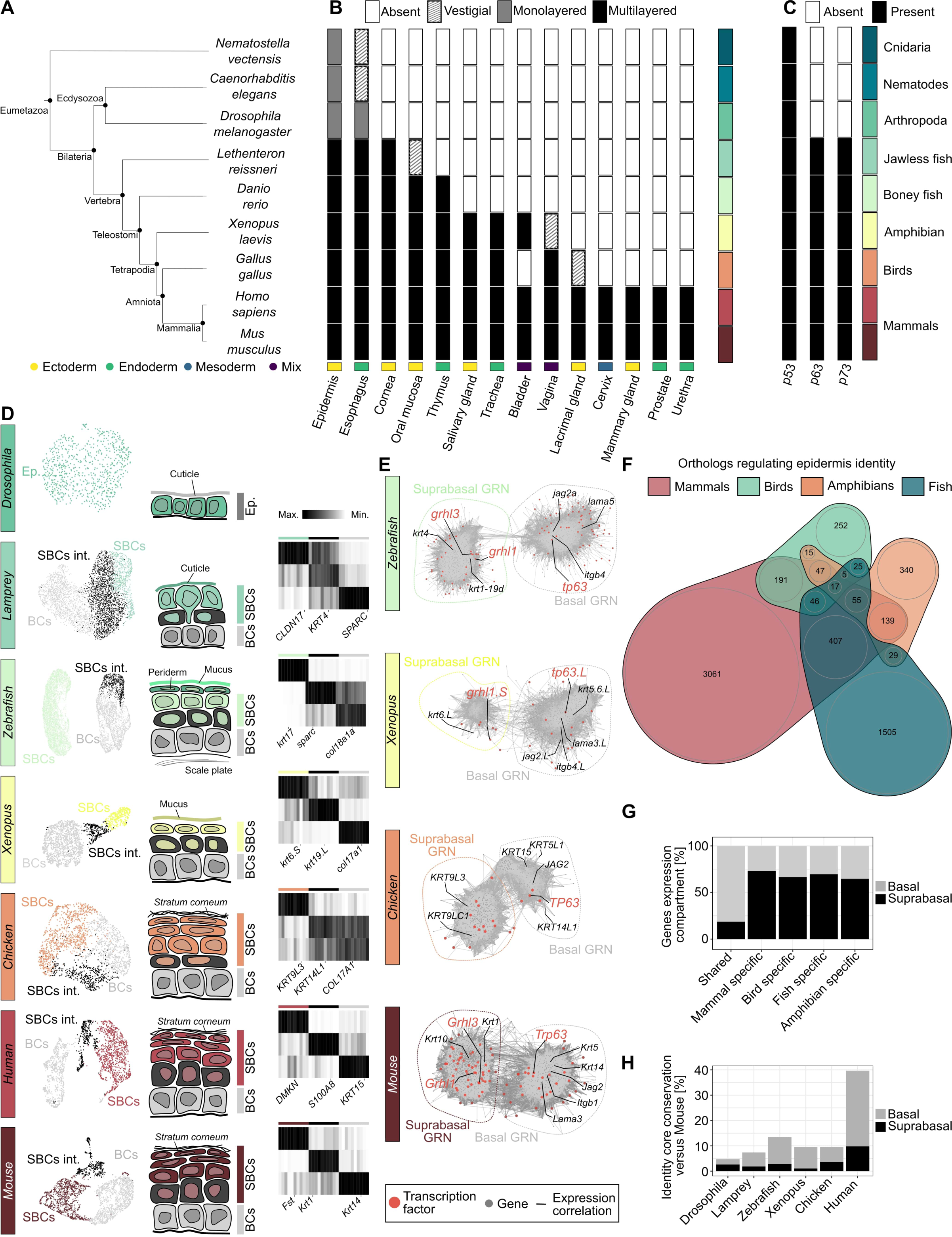
Identification of an ancestral GRN regulating epithelial stratification. ***A-B:*** *Phylogenetic tree (A) and the associated presence or absence of multilayered epithelia within each organism (B). Tissues are color-coded by the embryonic germ layer of origin. **C:** Presence or absence of genes of the p53 family within each organism. **D:** UMAP of scRNA-seq data colored by organ and cell cluster across Drosophila, Lamprey, Zebrafish, Xenopus, Chicken, mouse and Human epidermis. Associated scheme of the epidermis structure and marker of each cell population. Ep.: Epithelium, int.: intermediate. **E:** Gene co-expression network in each epidermis scRNA seq dataset. **F:** Venn diagram of the murine orthologs present in the co-expression network across species in epidermis. **G:** Percentage of genes expressed in basal or suprabasal compartment of the epidermis for shared or species-specific group. **H:** Proportion of murine orthologs from Basal and Suprabasal-core modules detected in the epidermal GRN of each species*.

Epithelial stratification first arose as a vertebrate innovation, transforming early protective barriers into specialized tissues with distinct physiological functions. While different vertebrate lineages may have followed distinct paths of diversification, here we focus on the mammalian lineage, taking its organs as a reference (Figure 6B). This transition from simple to multilayered epithelia coincided with the expansion of the p53 gene family and the neofunctionalization of p63^29,88^, a key regulator of basal identity (Figure 6C).

To identify innovations underlying this shift, we compared epidermal transcriptomes across species from *Drosophila melanogaster* to mammals^89–94^ (Figure 6D). Invertebrate epidermis was single-layered, whereas fish and amphibians developed stratified structures expressing simple epithelial keratins in suprabasal cells. Birds and mammals further elaborated this program with multi-step differentiation and cornification (Figure 6D)^95^.

Cross-species GRN analysis revealed asymmetric conservation between compartments (Figure 6E, F, S6A). Orthologs of mouse basal genes including *p63*, *Jag2*, and other basal regulators were broadly present and expressed in the basal epidermal cells across vertebrates (∼83%) (Figure 6E). In contrast, suprabasal GRNs showed greater lineage specificity (∼60–75%), reflecting independent innovations (Figure 6G; S6B). Some suprabasal regulators, such as *Grainyhead* family members, were preserved across lineages, but most effectors diversified with ecosystems: *keratins* in birds, *mucins* in amphibians, and cornification factors in mammals (Figure 6E; S6C). Comparative analysis of murine *Basal-* and *Suprabasal-core* confirmed that basal genes are more evolutionarily constrained, with ortholog recovery decreasing with phylogenetic distance (Figure 6H).

These analyses reveal that the basal compartment of vertebrate epidermis is deeply conserved, maintaining basal identity through a core p63–Notch program, the same axis we observed in murine multilayered epithelia. In contrast, suprabasal layers have diversified in each lineage, layering tissue- and biotope-specific modules onto this conserved basal scaffold. This modular architecture links a stable basal regulatory axis to lineage-specific effector programs, exemplifying how vertebrate multilayered epithelia combine structural integrity with adaptive functional specialization across organs and species.

### Spatial and temporal partitioning of ancestral and novel gene networks drives epithelial diversification

Building on this framework, we asked how evolutionary innovation is incorporated within multilayered epithelia. Although the basal program is ancient and conserved, suprabasal compartments exhibit extensive transcriptional variation across tissues and species. To examine how innovation arose within this conserved framework, we assessed the phylogenetic age of expressed genes across epithelia. Younger genes were consistently enriched in suprabasal compartments, independent of the overall transcriptome age index (TAI)^96^ of each tissue (Figure 7A; S7A–B), indicating that suprabasal layers function as evolutionary hotspots where novel modules can be recruited without disrupting core programs.

**Figure 7.**
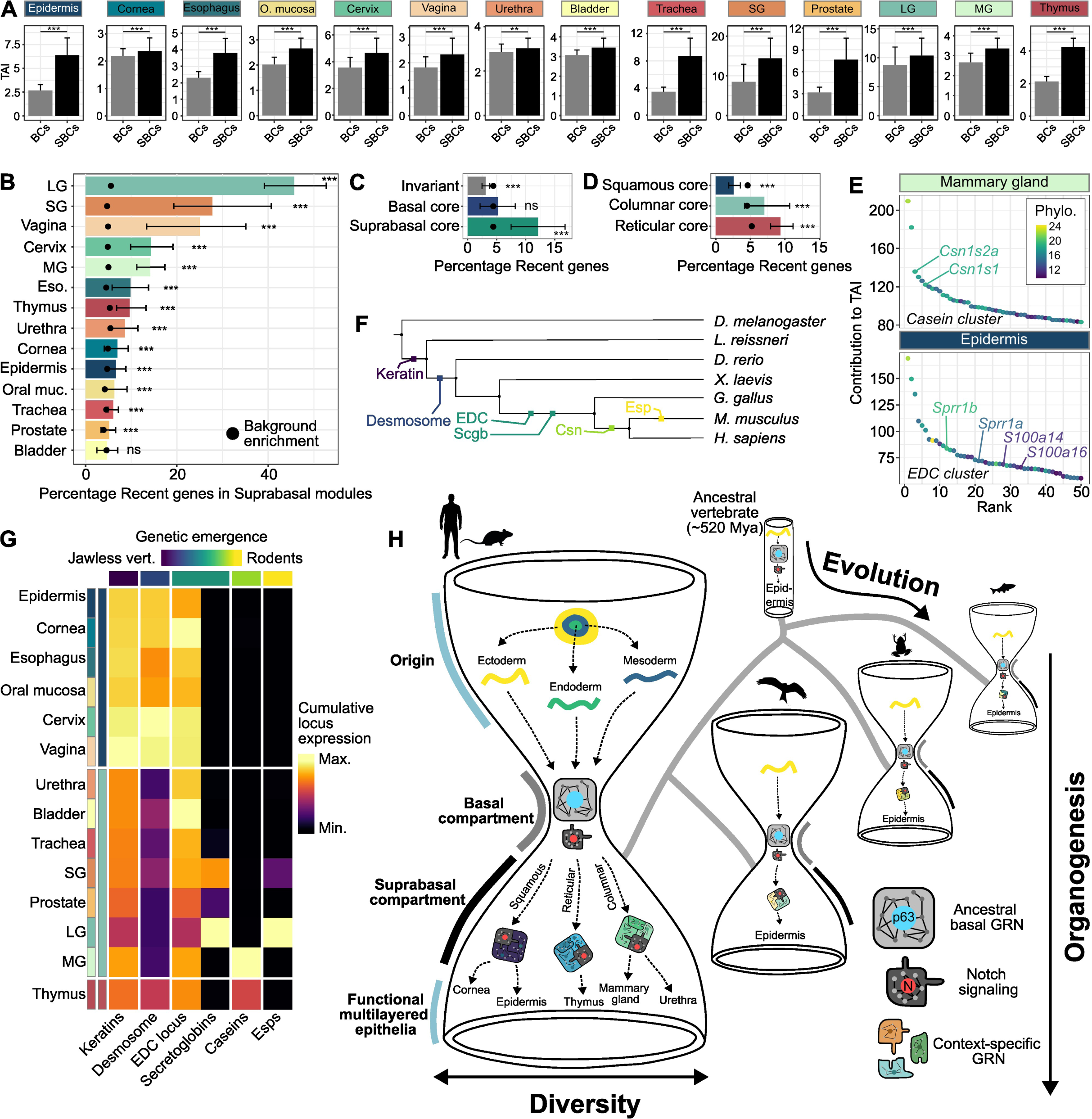
Evolution of tissue specific transcriptome through gene toolkit evolution and GRN rewiring. ***A:*** *Bootstrapped TAI calculated in pseudo bulk basal and suprabasal samples based on identity specific genes. Data displayed as mean + sd. Non parametric Wilcoxon test. **B**: Bootstrapped Enrichment in Recent genes (phylostratum >8) in tissue specific suprabasal modules compared to background enrichment in the same sample. Data displayed as mean + sd. Non parametric Wilcoxon test. **C**: Bootstrapped Enrichment in Recent genes in invariant and identity core modules compared to background enrichment in whole genome. Data displayed as mean + sd. Non parametric Wilcoxon test. **D**: Bootstrapped Enrichment in Recent genes in architecture specific modules compared to background enrichment in related sample. Data displayed as mean + sd. Non parametric Wilcoxon test. **E:** Main contributor to TAI in Mammary gland and Epidermis. Genes from casein and EDC complex are annotated respectively. **F:** Phylogenetic tree and associated genetic event related to emergence of each gene cluster. **G:** Cumulative expression of gene clusters in each murine multilayered epithelium. **H.** Developmental and evolutionary hourglass model of multilayered epithelia. During gastrulation, epithelia of distinct germ-layer origins (ectoderm, mesoderm, endoderm) emerge (top). Across vertebrates, these diverse embryonic inputs converge on a conserved basal bottleneck governed by the p63–Notch regulatory module (waist), representing a developmental and evolutionary constraint inherited from the ancestral vertebrate (∼520 Mya). From this shared basal program, lineage- and tissue-specific suprabasal GRNs (bottom) drive the diversification of multilayered epithelia into squamous, reticular, and columnar architectures*.

Analysis of our 14-tissues gene modules confirmed this trend. Recently evolved genes (phylostrata > 8) were enriched in suprabasal modules, up to sixfold in the lacrimal gland (Figure 7B; S7C). Consistent with this, the *Suprabasal-core* itself was consistently enriched (Figure 7C), whereas the Invariant module was depleted and the *Basal-core* showed no bias. Together, these differences reveal a hierarchical organization of evolutionary constraint: basal programs remain anchored in the ancestral GRN, while suprabasal modules are continually remodeled through assimilation of younger genes.

The same principle extended to morphological organization. The *Squamous-core*, representing the earliest stratified architecture, was dominated by conserved genes, whereas *Reticular* and *Columnar* cores, associated with later-evolving secretory, transitional and immunological epithelia, contained more recently evolved genes (Figure 7D). This evolutionary gradient mirrors the sequential emergence of epithelial specializations, with new gene modules progressively incorporated as tissue architectures diversified in the mammalian lineage.

Two representative gene clusters well illustrate this process. The epidermal differentiation complex (EDC), required for barrier formation, and the casein locus, essential milk components, both contribute disproportionately to the elevated TAI of epidermis and mammary gland (Figure 7E). Each represents a recently expanded locus that has become transcriptionally integrated into pre-existing epithelial GRNs. In the skin, EDC genes are targets of the ancestral regulators p63 and Grainyhead-like (*Grhl*) ^97–100^, indicating that newly acquired enhancers have linked this cluster to the ancient and conserved stratified-epithelium network^101–103^. In the mammary gland, casein genes, which emerged in early mammals, are co-regulated by broadly conserved factors such as the Notch pathway together with mammal-specific hormonal regulators including Stat5^104,105^, also reflecting the addition of new regulatory inputs onto an ancestral scaffold. These examples illustrate how recently evolved clusters are functionally assimilated through enhancer-level integration into existing GRNs, expanding tissue-specific repertoires while preserving overall network architecture.

To test the generalization of this principle, we compared six epithelial gene clusters: *keratins*, *desmosomal cadherins*, *EDC*, *secretoglobins*, *caseins*, and *exocrine-gland secreted peptides* (ESPs). Comparative genomics revealed a consistent trend: ancient clusters (keratins, desmosomal cadherins) are broadly expressed across epithelia, consistent with their conserved structural function, whereas recent clusters (caseins, ESPs) show restricted, lineage-specific expression (Figure 7F–G; S7D–E).

Collectively, these findings reveal that epithelial evolution is governed by a spatial and temporal partition of regulatory innovation. Basal layers preserve tissue integrity through an ancestral GRN of deeply conserved genes, whereas suprabasal layers incorporate newly emerged genes into dynamic, tissue-specific circuits. Within each epithelium, these ancient and recent programs coexist, coupling evolutionary continuity with adaptive change (Figure 7H). This hierarchical organization provides a unified framework for epithelial evolution, showing how tissues acquire new functions while maintaining a robust ancestral core.

## Discussion

Organogenesis of multilayered epithelia relies on a conserved transcriptional program that organizes epithelial cells into a characteristic layered architecture across vertebrate organs. The convergence of multilayered epithelia on this shared architecture raises the question of how this organization emerges across germ layers. The transcription factor p63, long known to orchestrate basal identity in ectodermal epithelia^14–16,106,107^, is first expressed during surface ectoderm specification^108^ and actively suppresses mesodermal and endodermal fates in vertebrate embryos^109–114^. Yet the repeated activation of p63-associated basal program in mesoderm- and endoderm-derivative to initiate tissues such as the esophagus, prostate, and cervix, shows that it can be assembled *de novo*, redeploying an ancestral ectodermal module outside its lineage of origin. These multiple heterotopies^115^, potentially triggered by shared stromal or placode-like inductive cues, illustrate how evolutionary innovation can arise through redeployment and of pre-existing regulatory circuits, a “toolkit” strategy as proposed by Jacob^116^.

While p63 provides a conserved entry point for establishing basal identity across germ layers, its functional output diverges across tissues. In rapidly renewing epithelia such as the epidermis, vagina, and esophagus, basal cells act as multipotent progenitors^117–119^, whereas in low-turnover glandular tissues they are mostly unipotent^120–124^. Yet, basal cells show limited diversification in gene expression compared with suprabasal compartments, suggesting that their tissue adaptation occurs primarily through modulation of cellular potency. Interestingly, the diversification of suprabasal gene expression may contribute to this process, as suprabasal ligands have been shown to signal to neighboring basal cells to restrict their potential^125^.

Suprabasal diversity emerges from the modular combination of Notch signaling with architecture- and tissue-specific transcriptional programs, generating columnar, squamous or reticular cells and their differentiation into secretory, cornified, or immunological units. How these modules are integrated with Notch remains largely unknown. Defining downstream effectors beyond HES1 is challenging, partly because Notch signaling operates as a multiprotein complex whose tissue-specific composition and cofactors dictate its activating or repressive functions^126,127^. Central open questions are whether Notch selectively engages tissue-specific targets, and to what extent local chromatin states shape its accessibility and regulatory output. Resolving these mechanisms will reveal how a single, conserved pathway can simultaneously coordinate fate commitment and functional diversification.

The ability of Notch signaling to integrate with distinct architectural and tissue-specific programs illustrates a broader evo-devo principle: gene regulatory networks act as combinatorial, reusable building blocks^128^. Yet, despite extensive morphological diversification, the multilayered epithelial regulatory program has remained deeply conserved for over half a billion years. This persistence likely stems from the essential developmental roles of its core factors, most notably p63, whose mutation disrupts limb, skin, and gland formation across vertebrates^14,15,106^. Multilayered organization may have resolved this constraint by spatially partitioning ancestral and derived regulatory modules, with new suprabasal programs built atop a conserved basal scaffold. This compartmentalized evolvability likely reflects dual selective pressures: developmental robustness preserving the ancestral GRN, and ecological adaptation driving diversification.

This study outlines a developmental, spatial, and evolutionary hourglass encoded in tissue architecture. Early epithelial progenitors from diverse germ layers converge on a conserved and developmentally constrained scaffold that dictate basal identity. From this constrained core, suprabasal programs radiate, integrating tissue- and architecture-specific modules shaped by environmental pressures. By unifying developmental and evolutionary perspectives, our work provides a conceptual framework to understand the emergence and diversification of multilayered epithelia.

## Resource availability

### Lead contact

Requests for further information and resources should be directed to and will be fulfilled by the lead contact, Robin P. Journot

### Material availability

This study did not generate new unique reagents.

### Data and code availability

No standardized data types were generated in this study. All data supporting the conclusions can be found in the main text or are available upon request from the corresponding authors, Robin Journot and Silvia Fre. The code used for the bioinformatic analysis will be available on Github upon publication. Any additional information required to reanalyze the data reported in this work paper is available from the corresponding authors upon request.

## Acknowledgments

We wish to acknowledge all Fre team members, as well as our colleagues Dr. Yohanns Bellaiche, Dr. Edouard Hannezo and Prof. Michel Labouesse, for technical advice and constructive discussions. The authors acknowledge the Cell and Tissue Imaging Platform PICT-IBiSA, member of the national infrastructure France-Bioimaging (https://ror.org/01y7vt929), supported by the French National Research Agency (ANR-24-INBS-0005 FBI BIOGEN) and the Cell and Tissue Imaging (PICT-IBiSA), Institut Curie, member of France-Bioimaging of the Genetics and Developmental Biology Unit (UMR3215/U934), as well as the In Vivo Experimental Facility, particularly Cedrick Pauchard, Mickael Garcia and Celine Daviaud, for help in the maintenance and care of our mouse colony.

This work was funded by, the French National Research Agency (ANR) grant numbers ANR-21-CE13-0047, ANR-22-CE13-0009 and ANR-24-CE13-3529-01, the Medical Research Foundation FRM "FRM Equipes" EQU201903007821, the FSER (Fondation Schlumberger pour l’éducation et la recherche) FSER20200211117, the Association for Research against Cancer (ARC) label ARCPGA2021120004232_4874, the Worldwide Cancer Research Foundation # 24-0216, the Ligue contre le Cancer label ELF_FRE_26738 and by Labex DEEP ANR-Number 11-LBX-0044.

R.J. was funded by a PhD fellowship from the French minister of research and by the FRM (FDT202404018370).

C.M. was funded by a postdoc fellowship from ARC (ARCPDF12021020003033) and Labex DEEP.

The funders had no role in study design, data collection and analysis, decision to publish, or preparation of the manuscript.

## Author Contributions

R.J., C.M. and S.F. conceived and designed the experiments. R.J., C.M., M.H. and E.K. performed all experiments. M.M.F. contributed to experimental work and edited the manuscript. R.J. performed all bioinformatic analysis. R.J. and C.M. curated, and interpreted results and prepared the figures. S.P. and M.S. provided critical remarks on the manuscript. R.J., C.M. and S.F. wrote the manuscript. S.F. provided funding and project administration. R.J. provided supervision. All authors reviewed and approved the manuscript.

## Methods

### Ethics statement

All studies and procedures involving animals were carried out in strict accordance with the recommendations of the European Community (2010/63/UE) for the protection of vertebrate species used for experimental and other scientific purposes. Approval was granted by the Ethics Committee of the Institut Curie CEEA-IC #2021-029 and the French Ministry of Research (reference #34364-202112151422480). We adhere to the internationally established principles of Replacement, Reduction and Refinement (3Rs) according to the Guide for the Care and Use of Laboratory Animals (NRC 2011). Animal husbandry, supply, maintenance and care in the animal facility of the Institut Curie (facility license #C75-05-18) before and during experiments fully met the needs and welfare of the animals. All animals were housed in individually ventilated cages under a 12:12h light/dark cycle, with water and food available ad libitum. All mice were culled by cervical dislocation. Mice were genotyped by PCR on genomic DNA extracted from an earpiece for adult mice or tail tip for embryos. Mouse breeding and husbandry was managed using the mouse colony organization software MiceManager: https://infenx.com/mouse-colony-management-software/.

### *In vivo* lineage tracing

We crossed the *K5-Cre^ERT2^* line^121^ with *mTmG*^84^ or *N1ICD-nGFP*^83^ for adult *in vivo* lineage tracing experiments. All mice used were in a mixed genetic background. Reporters and mutant alleles were recombined by a single intraperitoneal injection of tamoxifen free base prepared in sunflower oil containing 10% ethanol (0.1c:mg per g of mouse body weight), unless indicated otherwise.

### Immunostaining

Tissues samples were fixed in 4% PFA (2 hours at RT), washed (1h in PBS at RT) and incubated 48 hours in 30% sucrose at 4°C before embedding in optimal cutting temperature (OCT) compound. 8 μm thick cryosections (15 µm for the mammary gland) were cut using a cryostat (Leica CM1950) and stored at -20°C. For paraffin immunostaining, tissues were fixed overnight at 4°C in 4% PFA, washed 1 h in PBS and incubated in 70% ethanol until embedding. 5 μm thick sections were cut using a rotational microtome (Thermo scientific, Microm HM340E) and stored at RT. Section were deparaffinized, rehydrated and antigen retrieval was achieved by boiling slides in citrate buffer pH=6 for 15 minutes. Paraffin and frozen sections were blocked (PBS, 5% FBS, 1% BSA) for 1 hour and incubated with primary antibodies diluted in blocking buffer overnight in a humidified chamber. After washing, sections were incubated with a solution of secondary antibodies and DAPI (Merck) diluted in PBS for 1 hour. Finally, the sections were mounted on a slide (Aqua-Polymount, Polysciences). A list of all the antibodies used is provided in Table S1.

### Image acquisition

Images were acquired using a LSM780, LSM880 or LSM900 inverted laser scanning confocal microscope (Carl Zeiss) equipped with 25x/0.8 OIL LD LCI PL APO or 40x/1.3 OIL DICII PL APO objectives.

### Public transcriptomic data collection

Publicly available scRNAseq/ChIPseq datasets of the 14 murine adult tissues, and epidermis of metazoans were retrieved from the GEO database platform: https://www.ncbi.nlm.nih.gov/geo/ and public repository. All selected scRNA seq datasets were generated using the 10X technology and the expression matrix were retrieved from the processed output of standard pipeline. A list of all the datasets used can be found in Table S2.

### Statistics, reproducibility and data availability

Reproducibility codes for the analyses are available online via GitHub, as detailed in the ‘Code availability’ section. All statistical methods are described in the corresponding sections of the paper. Atlases of individual tissues and integrated dataset will be shared with associated metadata upon publication. Experiments were performed in biological replicates as indicated. A biological replicate corresponds to different animals. Experiments with at least 3 biological replicates were used to calculate the statistical value of each analysis. All graphs are mean + SD. Statistical analysis were performed using the Wilcoxon test or Chi squared test and are stated in figure legend. *: p-value < 0.05, **: p-value < 0.01, ***: p-value <0.001, ****: p-value < 0.0001.

### Transcriptomic dataset pre-processing

Individual datasets were pre-processed individually using Seurat^129^ to subset viable and good quality cell transcriptome based on the number, individual transcript detected per droplet and percentage of mitochondrial genes. As dataset originated from various sequencing platform and advance in the technology, the QC threshold were adjusted for each dataset. Globally, cells with too low *nFeatures_RNA*, too low or high *nCount_RNA* or percent.mt > 10 were discarded. In each dataset, low-quality genes were filtered out by excluding features detected in fewer than three cells with >1 UMI, cells with fewer than 500 total UMIs were also removed.

Gene expression counts were normalized using the *SCTransform*() function with UMI correction enabled and the top 3000 variable features retained. Principal Component Analysis (PCA) was performed on the SCT assay with 50 dimensions, and elbow plots were used to select relevant components for downstream analysis. UMAP embedding was computed using the accurate number of principal components, followed by neighborhood graph construction

### Cell type annotation

To annotate cell types, we first generated atlases of individual tissues through CSS integration (See Data integration section). The markers of each cluster were identified through differential expression and were compared with known markers. The analysis performed in the original publication of the datasets were qualitatively compared with our annotation to homogenize the process. The number of clusters was adjusted based on resolution of the *FindCluster()* function.

### Data integration

To construct a unified adult Multilayered epithelium atlas, we integrated annotated single-cell RNA-seq datasets from 14 distinct tissues: bladder, cervix, cornea, epidermis, esophagus, lacrimal gland, mammary gland, oral mucosa, prostate, salivary gland, thymus, trachea, urethra, and vagina. For each tissue, a previously annotated Seurat object was loaded and processed to harmonize assay formats and reduce memory usage.

To avoid barcode collisions during integration, cell names were prefixed with the tissue label using *RenameCells(*. Non-essential layers, including *scale.data* and *data* slots, were removed using regular-expression-based filtering (*removeLayersByPattern()*), and each object was reduced to its essential components using *DietSeurat()*.

Datasets were merged iteratively using Seurat’s *merge()* function to optimize memory handling. After each merge, the resulting object was joined using JoinLayers() to align assay layers across datasets. The merged object (Sample) thus represented a unified, layer-aware collection of epithelial cells from all tissues.

The integrated object was normalized using *NormalizeData()*, and the top 3,000 variable genes were identified using the variance-stabilizing transformation (*FindVariableFeatures(method = "vst")*).

To improve global alignment and overcome residual batch effects, we applied *Cluster Similarity Spectrum* (CSS) embedding using *cluster_sim_spectrum()* with Spearman correlation as similarity metric and kernel spectrum construction spectrum_type = ‘corr_kernel’) (“simspec”^72^ package). UMAP was recomputed in CSS space using the first 300 CSS components (*RunUMAP(reduction = "css", dims = 1:300)*), and refined clustering was performed using the first 200 CSS dimensions (*FindNeighbors(dims = 1:200), FindClusters(resolution = 0.1)*).

### Evaluation of batch effect correction across replicate

To evaluate the effectiveness of batch correction across epithelial biological replicates, we used the Local Inverse Simpson’s Index (LISI) as a quantitative metric for local batch mixing. LISI estimates the diversity of batch labels in each cell’s neighborhood, based on the inverse Simpson index of the k-nearest neighbor distribution (See section: *Measure of clustering diversity)*. High LISI values indicate that cells from different replicates are well intermixed, while low values (LISI ≈c:1) indicate that cells co-localize primarily with others from the same batch, suggesting residual batch effects.

To avoid conflating batch effects with true biological variation across tissues, LISI was computed within each tissue type. For each epithelial tissue, we subset the integrated Seurat object to retain only cells belonging to that tissue. From the corrected CSS embedding (first 30 dimensions), we extracted a low-dimensional matrix for LISI analysis. The *compute_lisi()* function from the lisi package^130^ was applied to this embedding, using the replicate label to estimate LISI per cell. For each tissue, we reported the mean LISI across all cells, which provides a robust summary of batch mixing quality. To confirm that improvements were due to the integration procedure, we repeated the same computation using the uncorrected PCA embedding, and compared before-versus-after mean LISI per tissue. Only tissues with at least two biological replicates and more than 50 cells were retained for evaluation. This approach allowed us to assess technical batch removal in a biologically meaningful way, preserving within-tissue variability while excluding inter-tissue differences.

### Identification of differentially expressed genes and tissue specific markers

Differentially expressed genes (DEGs) were identified using the *FindMarkers()* function in Seurat, applying the Wilcoxon rank-sum test to compare each epithelial tissue against all others. Only genes expressed in at least 10% of cells in either group and with an average log fold change ≥ 0.25 were retained. P-values were adjusted using the Bonferroni method, and genes with an adjusted p-value < 0.01 were considered significant. The top genes were selected for representation as indicated in figure legends. To define tissue-specific genes, we selected DEGs that were significantly upregulated in one tissue compared to all others and showed minimal or no expression in the remaining tissues (expression in < 10% of cells across other tissues).

### Pseudo bulk expression and cosine similarity analysis of cell class

To assess transcriptomic similarity between cell class cells across different Multilayered epithelial tissues, we performed a pseudo bulk expression analysis followed by cosine similarity and hierarchical clustering. Pseudo bulk expression matrices were generated for immune cells using the *AverageExpression()* function in Seurat on a down sampled dataset to a maximum of 1,000 cells per tissue–cell type combination. The resulting matrix was converted to a standard numeric matrix, and cosine similarity between all tissue-level immune cell profiles was computed using the *proxy::simil()* function with the "cosine" method.

The cosine similarity matrix was converted into a distance matrix (Euclidean distance of cosine values) and subjected to hierarchical clustering using the *hclust()* function with the "ward.D2" method.

### Measure of clustering diversity

Local Inverse Simpson’s Index (LISI) is a metric to quantify local diversity in a low-dimensional embedding like UMAP used in single-cell data analysis to assess cell types are mixed.

Local Inverse Simpson’s Index (LISI) was computed to quantify the local diversity of cell classes (e.g., embryonic origin or epithelia architecture) in the integrated UMAP space. For each cell, LISI was calculated based on the label distribution among its k nearest neighbors (typically k = 30).

For each cell, LISI was computed as:

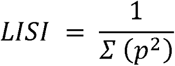

where p□: is the proportion of neighbors belonging to category i. The score reflects the effective number of distinct categories in the neighborhood:

LISI = 1 indicates that all neighbors belong to a single category (no diversity).

LISI = N (where N is the number of categories) indicates an equal distribution of neighbors across all categories (maximum diversity).

### KEGG pathway enrichment

Basal GRN genes were converted from mouse gene symbols to Entrez identifiers using the *bitr()* function from clusterProfiler^131^. KEGG pathway enrichment was then performed with *compareCluster(fun = "enrichKEGG")* using the *Mus musculus* annotation, applying Benjamini–Hochberg correction (p < 0.05; q < 0.1). Pairwise semantic similarity between enriched terms was computed with *pairwise_termsim()* to allow redundancy assessment.

### Linear model of gene expression and module assignment

#### Preprocessing

Raw counts were aggregated into pseudo bulk profiles for each unique combination of tissue, cell identity, and cell type. Lowly expressed genes (counts per million < 2 in fewer than 5% of samples) and non-informative features (e.g., mitochondrial genes) were excluded. Normalization was performed using the trimmed mean of M-values (TMM) method to correct for differences in library size.

#### Cell-Means Model

The expression of each gene g in sample s was modeled using a cell-means approach, where the observed expression is represented as the sum of a global mean and a coefficient specific to the combination of Architecture (A), Tissue (T), and Identity (I) for that sample:

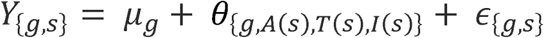

Here, *y_{g,s}_* is the expression of gene *g* in sample *s*, *µ_g_* is the baseline expression level for gene *g, B_{g,A(s),T(s),I(s)}_* is the deviation parameter for the specific combination of Function, Tissue, and Identity, and *c_{g,s}_* is the residual error. This model assigns a unique coefficient to each Function × Tissue × Identity combination, capturing the mean expression level for that group.

#### Contrast Decomposition

To interpret the cell-means coefficients biologically, we decomposed each deviation parameter β_{g,f,t,I}_ into a sum of contribution parameters using linear contrasts:

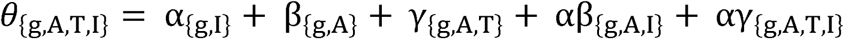

where:

- α_{g,I}_ global identity effect, capturing basal vs. suprabasal differences across all architectures and tissues
- β_{g,A}_ architectural effect, reflecting the average influence of each epithelial class (squamous, columnar, reticular)
- γ_{g,A,T}_ tissue effect nested within architecture, representing tissue-specific deviations relative to the mean of their architectural class
- αβ_{g,A,I}_ modulation of identity by architecture, describing how basal–suprabasal differences vary across epithelial architectures
- αγ_{g,A,T,I}_ modulation of identity by tissue, capturing tissue-specific adjustments of basal–suprabasal differences within each tissue architecture

Each component was isolated by applying orthogonal contrast matrices to the fitted coefficients, allowing independent estimation of variance attributable to each biological factor.

#### Module Assignment

Genes were assigned to functional modules based on the dominant source of variation in their expression, determined by statistical significance (adjusted p-value) and effect size (partial eta-squared, η^2^_p_) with:

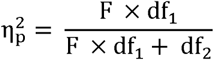

where F is the F-statistic, anddf_l_ and df_2_ are the degrees of freedom.

Genes were assigned to modules according to a hierarchical decision tree:

- Identity modules: if α_{g,I}_ was significant in ≥75% of tissues (adjusted p < 1e−10) and had the largest η^2^_p_. Genes were labeled Basal or Suprabasal depending on the direction of the effect.
- Architecture modules: if β_{g,A}_ was dominant (adjusted p < 1e-6, highest η^2^_p_).
- Tissue modules: if γ_{g,A,T}_ was dominant (adjusted p < 1e-6).
- Architecture × Identity modules: if αβ_{g,A,I}_ was significant (p < 1e−4) and dominant.
- Tissue × Identity modules: if αγ_{g,A,T,I}_ was significant (p < 0.1) and dominant.
- Invariant genes: if no effect passed thresholds.

This hierarchical assignment ensured that each gene was associated with the main factor explaining its variance in expression, providing a biologically interpretable partitioning of transcriptomic regulation across epithelial architectures and identities.

#### Validation of Model Assumptions

To ensure the validity of the linear model, we verified the following assumptions:

- Normality of Residuals:

The normality of residuals was assessed using Q-Q plots, which showed approximate normality with minor deviations at the tails.

- Homoscedasticity:

Residuals vs. fitted values plots were examined to confirm that the residuals were randomly scattered around zero, with no clear patterns, indicating homoscedasticity.

- Heteroscedasticity:

The relationship between the standard deviation of residuals and the mean fitted values exhibited a funnel shape, consistent with RNA-seq data. This heteroscedasticity was addressed by the precision weights applied during the voom transformation, which stabilizes variance across genes.

#### Implementation

All analyses were performed in R (v4.2.0) using *edgeR* for normalization, *limma* for voom transformation, model fitting, and empirical Bayes moderation.

### Permutation test for variance decomposition

To assess whether the observed variance decomposition reflected genuine biological modularity rather than random structure, we implemented a permutation test on the pseudobulk expression profiles. For each gene, partial η² values were estimated for the main effects of Identity 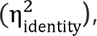 Architecture 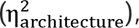 Tissue 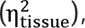 Architecture × Identity 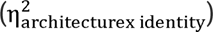 and Tissue × Identity 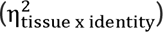 using linear modelling with *limma/voom* and custom contrast matrices.

We then computed the mean η² across all genes for each variance component. To generate a null distribution, we randomly permuted Tissue and Identity labels across pseudo bulk samples while keeping the expression matrix unchanged, rebuilt the design matrix, and repeated the variance decomposition. This procedure was iterated 500 times, yielding a distribution of mean η² values expected under random labeling. Permutation p-values were calculated as:

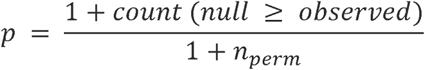

Observed values were compared against the null distributions using boxplots with the empirical value indicated as a black point.

### Cell-level variance attribution analysis

To extend gene-level variance partitioning to the single-cell scale, we developed an expression-weighted η² scoring approach. For each gene g, variance explained by identity, function, tissue, or interaction terms was estimated as η^2^ using the linear models described previously. To obtain cell-level values, we weighted each gene’s η² contribution by its normalized expression E*_{i,g}_* in cell i, and then computed the weighted mean across all expressed genes:

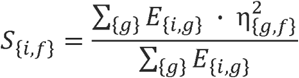

where S*_{i,f}_* is the variance attribution score of cells i for factor f. These yields, for each cell, a profile of scores reflecting the relative influence of identity-, function-, and tissue-level effects on its transcriptome.

### Gene co-expression inference

Raw UMI count data were extracted from the RNA assay of the Seurat object using the LayerData() function. To identify genes suitable for co-expression analysis, we computed, for each gene, the average expression across all cells in which the gene was detected (i.e., non-zero counts) as well as the percentage of cells expressing the gene. Genes with an average expression >1 in expressing cells and detected in >10% of cells were retained for downstream analysis.

Co-expression analysis was performed using the CS-CORE algorithm^132^ on each tissues, restricted to cells in the resting (non-proliferative) state. For each tissue, the CS-CORE function was run using the filtered gene list as input, and pairwise co-expression estimates and associated p-values were computed. Multiple testing correction was applied using the Benjamini–Hochberg procedure. Adjusted p-values were calculated for all gene pairs, but no filtering threshold was applied at this stage.

### Module activity inference by sparse regression

Raw counts from epithelial atlas were aggregated into pseudobulk profiles per tissue–identity group. Genes with CPM > 2 in at least 5% of groups were retained, normalized by TMM (*edgeR*), and transformed to log2-CPM with precision weights (*voom*, *limma*). Expression values were z-scored per gene across tissues. Gene modules were defined from the global module table and restricted to sets with >5 expressed genes. For each tissue pseudobulk, gene expression was decomposed into weighted module activities using lasso regression with non-negative coefficients (*glmnet*), applied to a design matrix containing positive and negative copies of each module to allow signed contributions. Final weights were collapsed into signed scores, z-scored across tissues, and displayed as a heatmap, with columns split by tissue function and rows split into invariant, functional core, tissue-core, and tissue-identity modules.

### GRN representation

To infer transcriptional GRNs across species, we combined co-expression analysis with differential expression data from single-cell RNA-seq of epithelial tissues in human, mouse, chicken, zebrafish, Xenopus, lamprey, and Drosophila. For each species, transcription factors (TFs) were retrieved from species-specific TF reference databases.

GRNs were constructed by filtering a species-specific co-expression matrix (see Gene co-expression inference) to retain only genes significantly differentially expressed genes between basal and suprabasal cells. Co-expression scores were threshold to generate adjacency matrices, and too weak links were removed. Resulting networks were defined as undirected graphs with genes as nodes and significant co-expression links as edges, weighted by co-expression strength. Networks were exported in GraphML format and visualized in *Cytoscape* using the Prefuse Force Directed layout.

### ChIPseq data visualization for p63 binding sites

To visualize p63 ChIP-seq signal across specific genomic regions, we used the R packages Gviz, rtracklayer, and GenomicRanges. The processed ChIP-seq signal for p63 (GSM2559737) was obtained in bedGraph format. Genomic regions of interest were defined using *GRanges* objects with *mm10* genome coordinates.

To display gene annotations, we used the *GeneRegionTrack* function from the *Gviz* package, built from the *TxDb.Mmusculus.UCSC.mm10.knownGene* annotation. A DataTrack object was created to plot ChIP-seq signal intensity as a histogram over the selected region.

All tracks, including the gene track, ChIP signal track, genome axis, and ideogram, were plotted using the plotTracks() function. Genomic regions were selected based on loci of interest, such as candidate target genes or regulatory elements.

### Quantification of promoter⍰centered p63 binding

GenomelZlwide p63 ChIPlZlseq coverage was computed from bedGraph tracks using rtracklayer and converted to a strandlZlindependent RleList capped at the 99.9th percentile to limit outliers. Gene coordinates were obtained from *TxDb.Mmusculus.UCSC.mm10.knownGene*, and promoter regions were defined as ±1c:kb around the transcription start site (TSS). For each promoter, the mean normalized ChIPlZlseq signal was extracted using the *ScoreMatrixBin()* function from the genomation package (bin.numc:=c:1). Promoters with mean signal above the 95th percentile of all promoters were considered “highly bound.” The fraction of highly bound promoters was computed for basallZlcore and background gene sets, and finally compared statistically using Fisher’s exact test.

### TSS⍰centered metagene profiles

To visualize binding profiles, the same coverage data were averaged in 50c:bp bins within ±2c:kb of each TSS using a custom extraction function. Mean signal andc:±c:SEM were plotted for basallZlcore and background promoters.

### Transcription factor motif enrichment analysis

To identify candidate transcriptional regulators associated with architecture-specific gene modules, we performed TF enrichment analysis using the ChIP-X Enrichment Analysis (ChEA3) database. For each architectural module (*Squamous-, Reticular-,* and *Columnar-core*), we provided the list of module genes as input. Enrichment was computed against integrated ChIP-seq reference datasets combining ENCODE, ReMap, and literature-curated TF–target interactions. TFs were ranked by their combined enrichment score, which integrates mean rank across multiple evidence types. The top enriched TFs were visualized as ranked enrichment plots, and representative regulators for each architecture were annotated in the figure. Colored annotated TFs correspond to genes that belongs to the module.

### Evolutionary Age Profiling and Tissue Transcriptome Analysis

Gene orthology and evolutionary age were obtained from the *phylomapr* package, which provides a Phylostratum annotation for Mus musculus genes from Barrera-Redondo et al. 2023^133^. UniProt identifiers were mapped to Ensembl gene IDs and gene symbols using the *biomaRt*^134^ package.

#### Transcriptome Age Index (TAI) Calculation Across Tissues

To assess the evolutionary age of gene expression profiles in adult epithelia, we computed the Transcriptome Age Index (TAI) per tissue. Raw gene expression counts were extracted from the Seurat object, and pseudobulk transcriptomes were generated by summing raw counts across all cells from each tissue. Normalization was performed using *log1p* transformation.

Genes were intersected with the phylogenetic map to retain those with known phylostrata. A *PhyloExpressionSet* was then constructed with rows representing genes and columns representing phylostratum and normalized expression. The TAI was calculated for each tissue using the *TAI()* function from the *myTAI* package^96^. The index reflects the average evolutionary age of expressed genes, weighted by their expression level.

#### Identification of Evolutionarily Biased Contributors

To identify genes contributing disproportionately to the TAI in each tissue, we computed a contribution score as the product of gene expression and phylostratum value (i.e., older genes contribute less).

#### Differential Expression and TAI in Cell Types

To compare evolutionary biases between basal and suprabasal compartments, we performed tissue-wise differential expression (DE) analysis. For each tissue containing both basal and suprabasal populations, marker gene detection was carried out with *FindMarkers()* using thresholds of logFC > 0.5 and minimum detection fraction > 0.1. The top 500 DE genes upregulated in each identity (basal or suprabasal) were retained.

For each tissue-compartment pair, pseudobulk transcriptomes were generated by summing raw gene counts across all cells, and normalized by dividing by the number of contributing cells to control for sample size. Genes were annotated with their phylostratum and the TAI was calculated as previously described. Briefly, TAI was defined as the weighted mean phylostratum of expressed genes, where gene weights correspond to normalized expression levels. Samples with fewer than 30 annotated genes were excluded.

To account for sampling variability, we applied a bootstrap strategy. For each pseudobulk sample, the set of DE genes was resampled with replacement 150 times, and TAI was recomputed in each replicate. This produced a bootstrap distribution of TAI values per tissue and identity. From these distributions, we estimated mean TAI, standard deviation, and 95% confidence intervals. Differences between basal and suprabasal bootstrap distributions were assessed using Wilcoxon rank-sum tests. Multiple testing correction across tissues was performed using the Benjamini–Hochberg method.

#### Venn Diagram Analysis

For exploratory purposes, Venn diagrams were generated to visualize overlap in top evolutionary contributors across tissues using *nVennR*. Gene sets were grouped by tissue and displayed interactively, with gene region intersections extracted for interpretation.

#### Phylostratum enrichment of gene modules

To assess whether specific gene modules were enriched in ancient or recently evolved genes, we annotated each gene with its phylostratum. Genes were binned into three evolutionary age classes: Very Ancient (Phylostratum ≤2), Intermediate (3≤ Phylostratum ≤8), and Recent (Phylostratum >8). Each module, defined as the set of genes assigned to a given identity/function/tissue component, was intersected with these age classes.

For each module and age class, enrichment was evaluated relative to a background set of expressed genes matched to the same tissue and identity context. Genes were considered expressed if they were detected in at least 25% of cells from the corresponding. Enrichment was tested by constructing 2×2 contingency tables of module vs. background membership and applying one-sided Fisher’s exact tests. To assess robustness, module fractions were compared against bootstrap resampling distributions (n=100) drawn from the same gene set, and significance was additionally tested with one-sample Wilcoxon tests against the background fraction.

## Supplementary Information

**Table S1:**
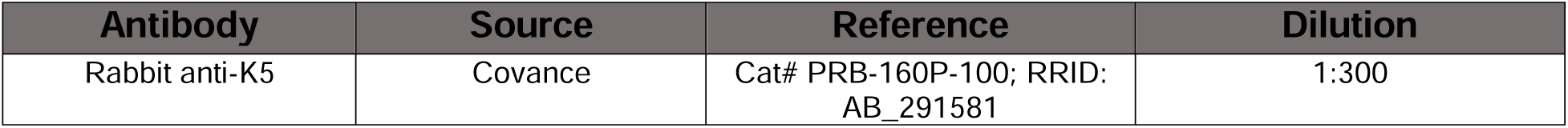
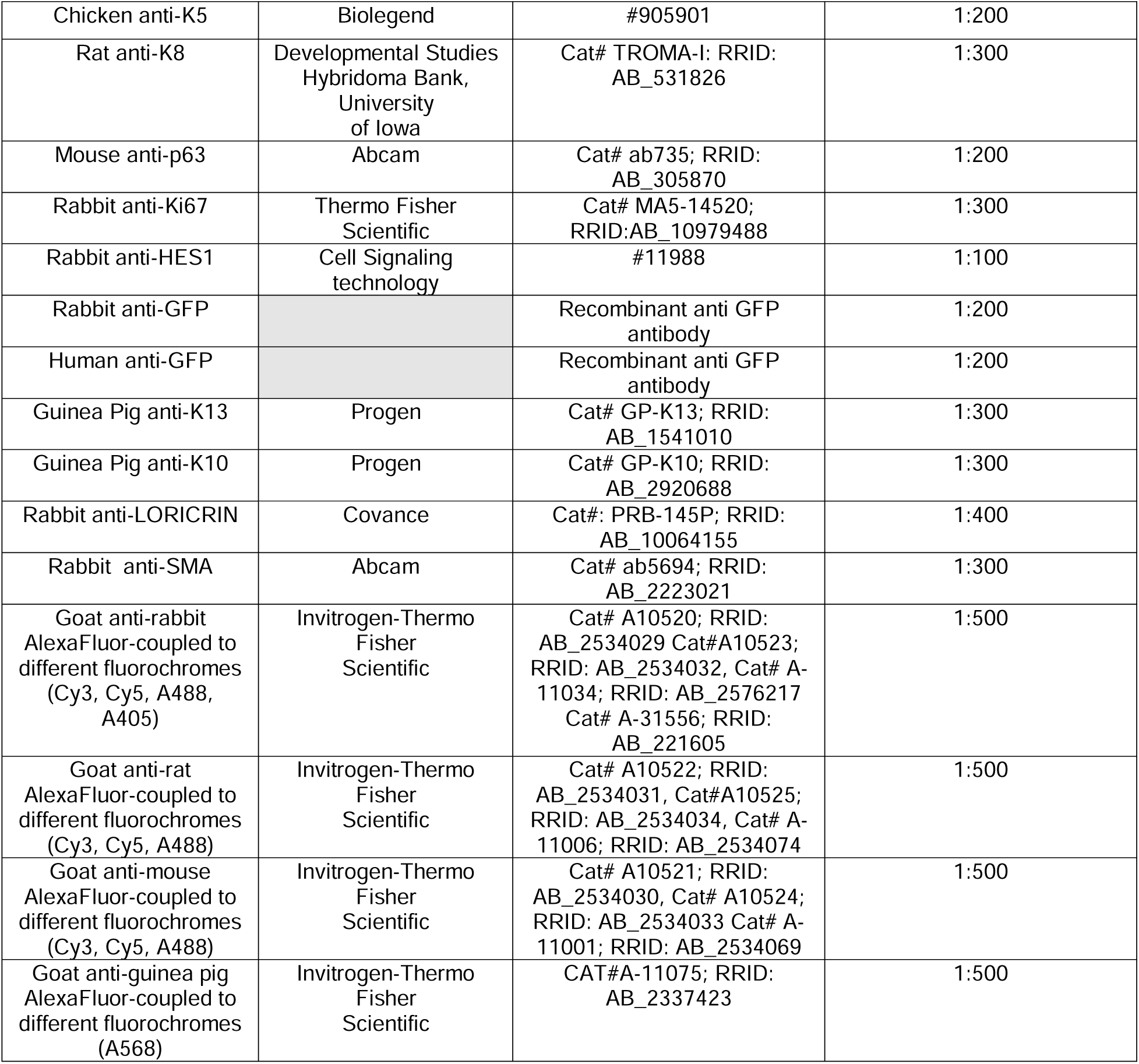
Antibodies.

| Antibody | Source | Reference | Dilution |
| --- | --- | --- | --- |
| Rabbit anti-K5 | Covance | Cat# PRB-160P-100; RRID: AB_291581 | 1:300 |
| Chicken anti-K5 | Biologend | #905901 | 1:200 |
| Rat anti-K8 | Developmental Studies Hybridoma Bank, University of Iowa | Cat# TROMA-I; RRID: AB_531826 | 1:300 |
| Mouse anti-p63 | Abcam | Cat# ab735; RRID: AB_305870 | 1:200 |
| Rabbit anti-Ki67 | Thermo Fisher Scientific | Cat# MA5-14520; RRID:AB_10979488 | 1:300 |
| Rabbit anti-HES1 | Cell Signaling technology | #11988 | 1:100 |
| Rabbit anti-GFP |  | Recombinant anti GFP antibody | 1:200 |
| Human anti-GFP |  | Recombinant anti GFP antibody | 1:200 |
| Guinea Pig anti-K13 | Progen | Cat# GP-K13; RRID: AB_1541010 | 1:300 |
| Guinea Pig anti-K10 | Progen | Cat# GP-K10; RRID: AB_2920688 | 1:300 |
| Rabbit anti-LORICRIN | Covance | Cat#: PRB-145P; RRID: AB_10064155 | 1:400 |
| Rabbit anti-SMA | Abcam | Cat# ab5694; RRID: AB_2223021 | 1:300 |
| Goat anti-rabbit AlexaFluor-coupled to different fluorochromes (Cy3, Cy5, A488, A405) | Invitrogen-Thermo Fisher Scientific | Cat# A10520; RRID: AB_2534029 Cat#A10523; RRID: AB_2534032, Cat# A-11034; RRID: AB_2576217 Cat# A-31556; RRID: AB_221605 | 1:500 |
| Goat anti-rat AlexaFluor-coupled to different fluorochromes (Cy3, Cy5, A488) | Invitrogen-Thermo Fisher Scientific | Cat# A10522; RRID: AB_2534031, Cat#A10525; RRID: AB_2534034, Cat# A-11006; RRID: AB_2534074 | 1:500 |
| Goat anti-mouse AlexaFluor-coupled to different fluorochromes (Cy3, Cy5, A488) | Invitrogen-Thermo Fisher Scientific | Cat# A10521; RRID: AB_2534030, Cat# A10524; RRID: AB_2534033 Cat# A-11001; RRID: AB_2534069 | 1:500 |
| Goat anti-guinea pig AlexaFluor-coupled to different fluorochromes (A568) | Invitrogen-Thermo Fisher Scientific | CAT#A-11075; RRID: AB_2337423 | 1:500 |

**Table S2:**
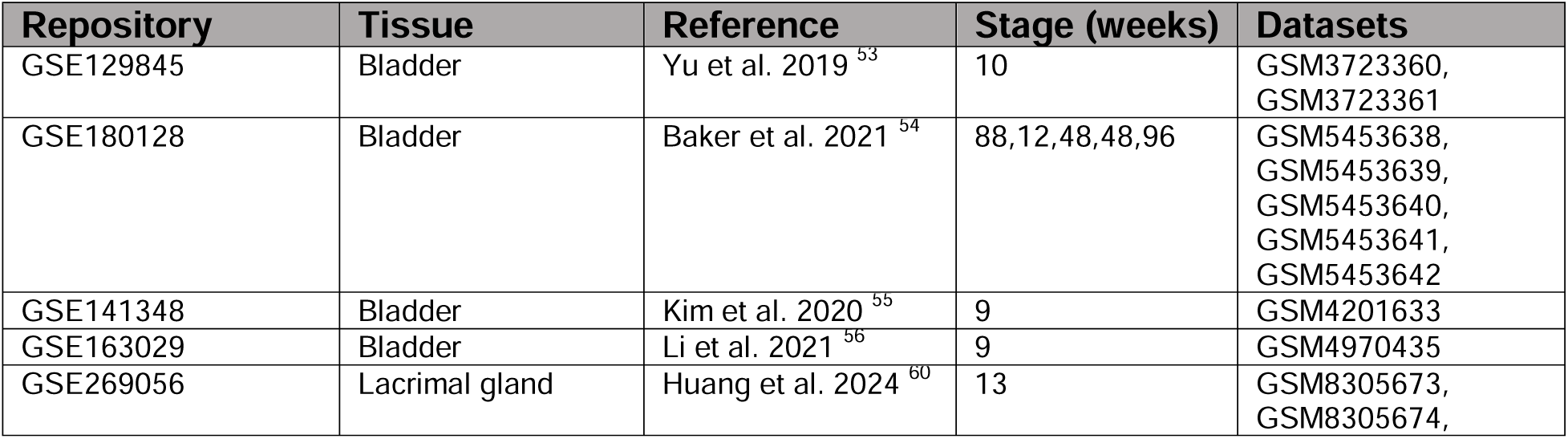
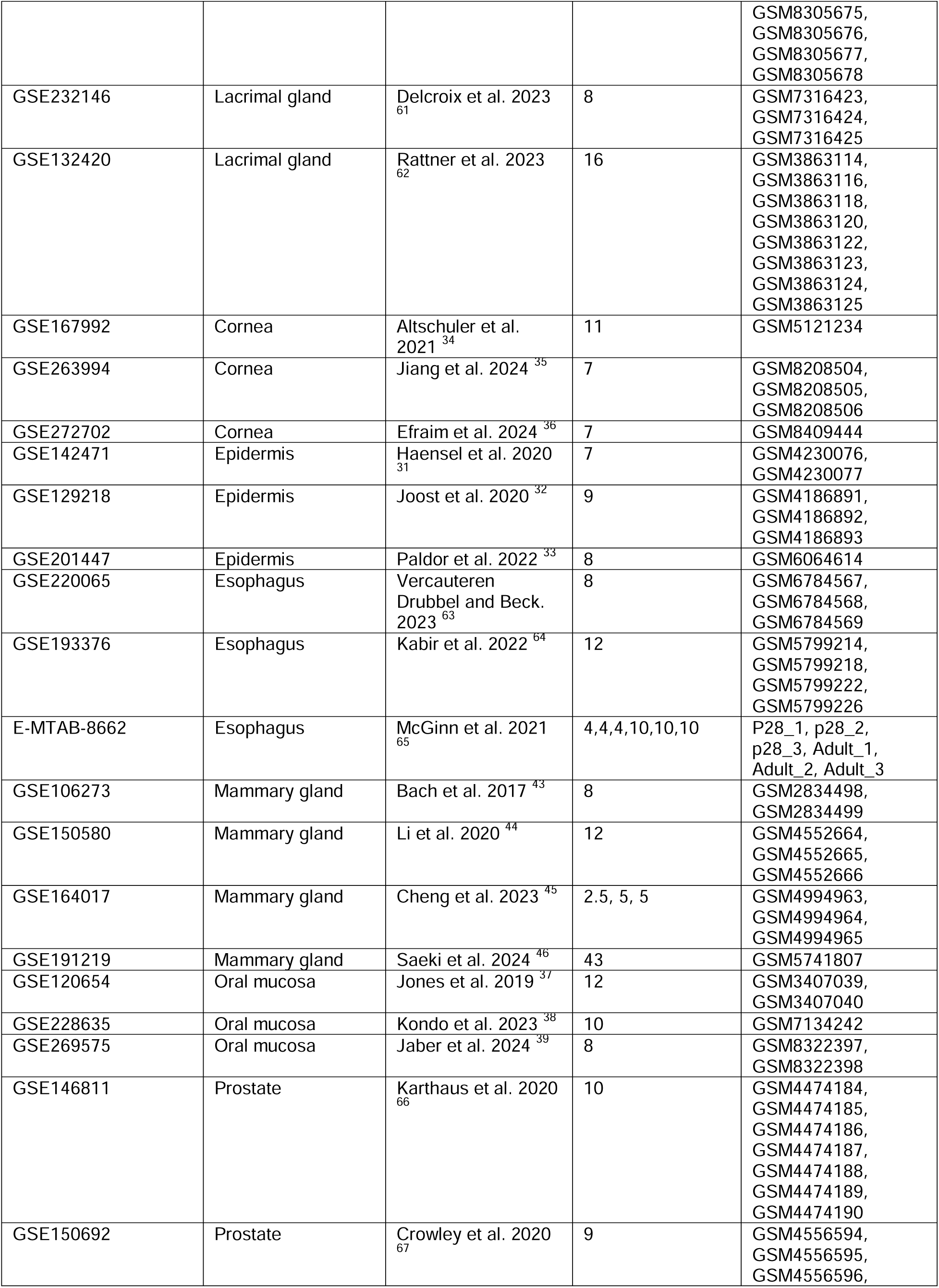
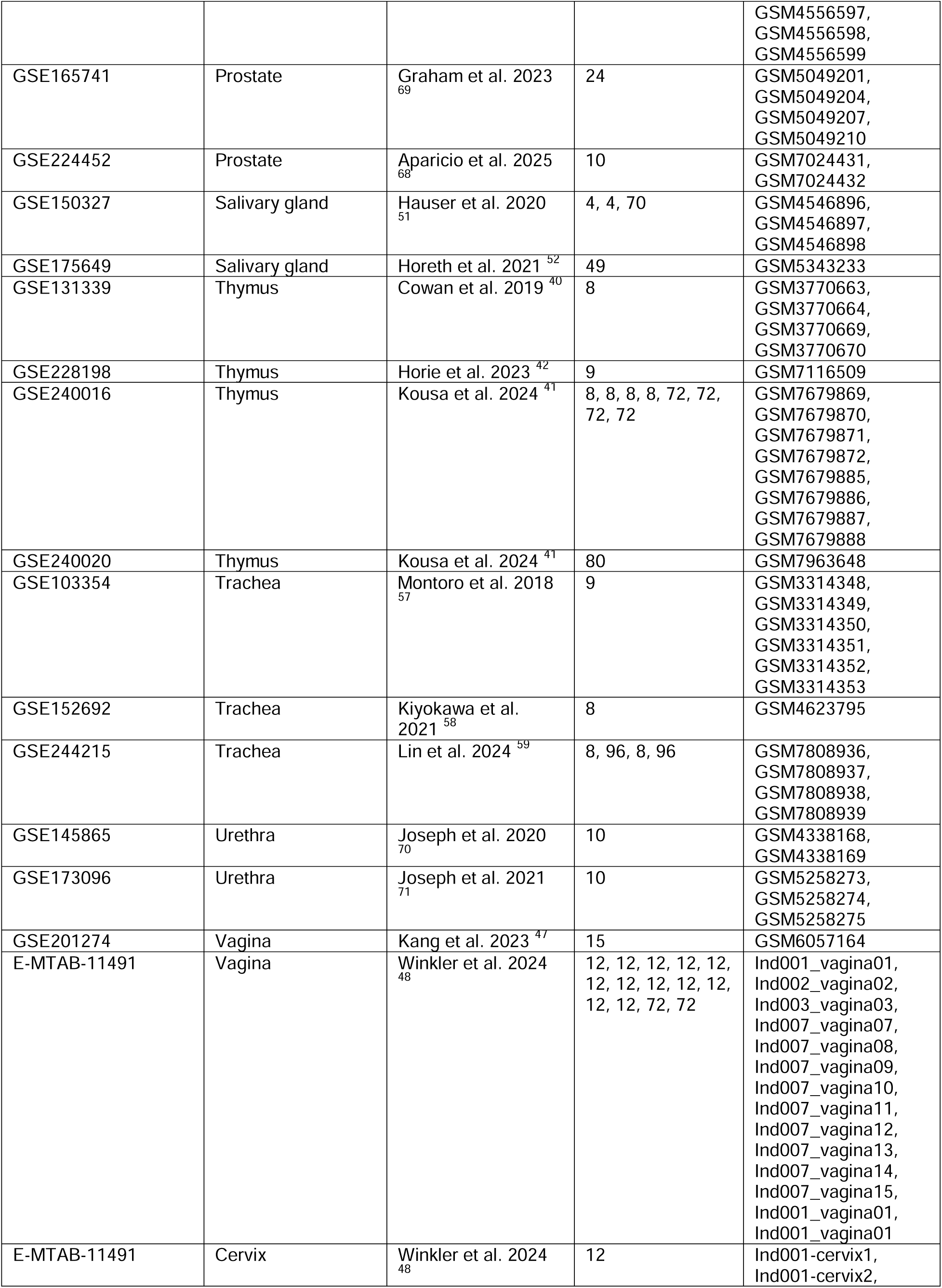
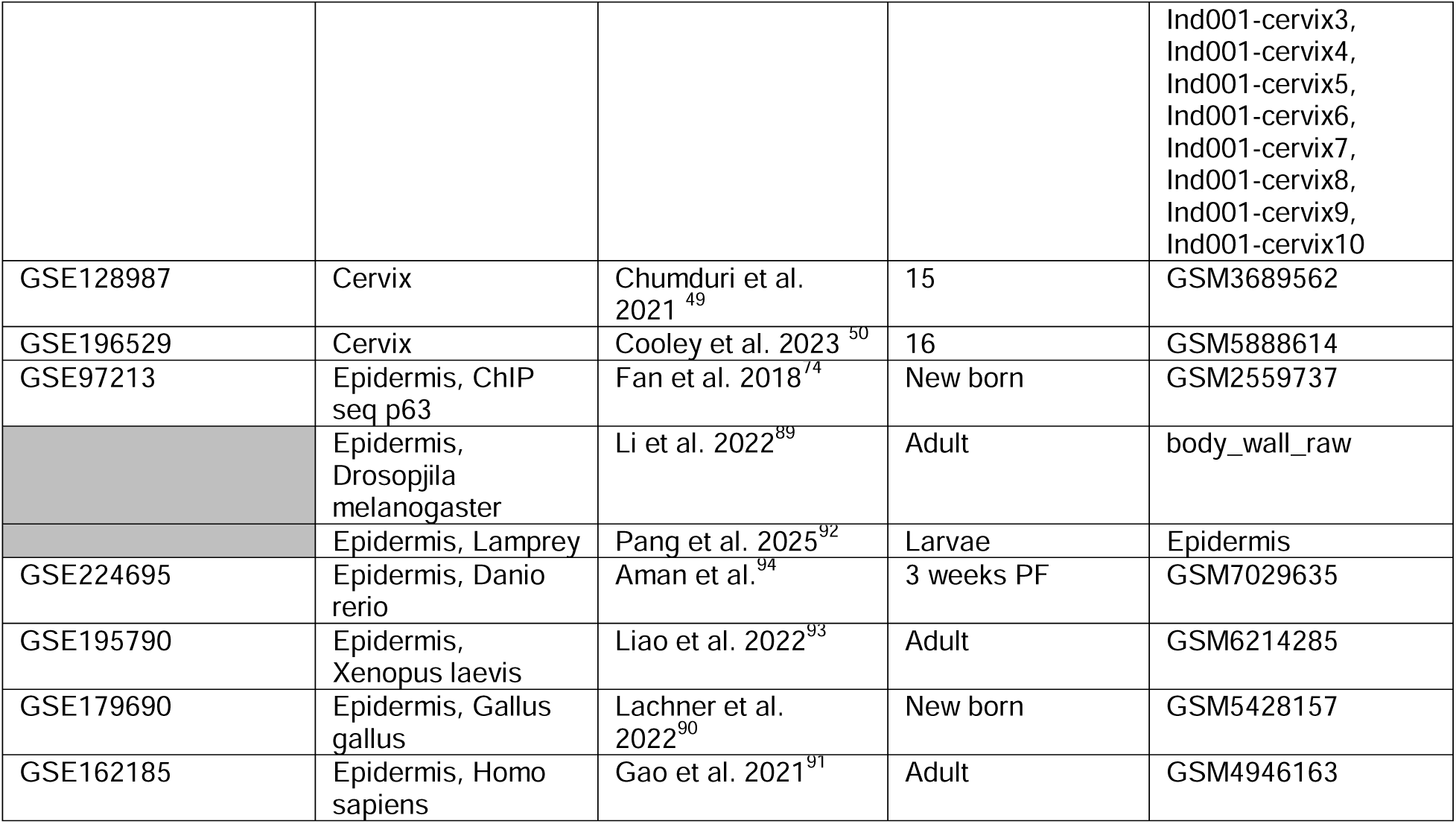
Public transcriptomic dataset.

| Repository | Tissue | Reference | Stage (weeks) | Datasets |
| --- | --- | --- | --- | --- |
| GSE129845 | Bladder | Yu et al. 2019<sup>53</sup> | 10 | GSM3723360, GSM3723361 |
| GSE180128 | Bladder | Baker et al. 2021<sup>54</sup> | 88,12,48,48,96 | GSM5453638, GSM5453639, GSM5453640, GSM5453641, GSM5453642 |
| GSE141348 | Bladder | Kim et al. 2020<sup>55</sup> | 9 | GSM4201633 |
| GSE163029 | Bladder | Li et al. 2021<sup>56</sup> | 9 | GSM4970435 |
| GSE269056 | Lacrimal gland | Huang et al. 2024<sup>60</sup> | 13 | GSM8305673, GSM8305674, |
|  |  |  |  | GSM8305675,<br>GSM8305676,<br>GSM8305677,<br>GSM8305678 |
| GSE232146 | Lacrimal gland | Delcroix et al. 2023<sup>61</sup> | 8 | GSM7316423,<br>GSM7316424,<br>GSM7316425 |
| GSE132420 | Lacrimal gland | Rattner et al. 2023<sup>62</sup> | 16 | GSM3863114,<br>GSM3863116,<br>GSM3863118,<br>GSM3863120,<br>GSM3863122,<br>GSM3863123,<br>GSM3863124,<br>GSM3863125 |
| GSE167992 | Cornea | Altschuler et al. 2021<sup>34</sup> | 11 | GSM5121234 |
| GSE263994 | Cornea | Jiang et al. 2024<sup>35</sup> | 7 | GSM8208504,<br>GSM8208505,<br>GSM8208506 |
| GSE272702 | Cornea | Efraim et al. 2024<sup>36</sup> | 7 | GSM8409444 |
| GSE142471 | Epidermis | Haensel et al. 2020<sup>31</sup> | 7 | GSM4230076,<br>GSM4230077 |
| GSE129218 | Epidermis | Joost et al. 2020<sup>32</sup> | 9 | GSM4186891,<br>GSM4186892,<br>GSM4186893 |
| GSE201447 | Epidermis | Paldor et al. 2022<sup>33</sup> | 8 | GSM6064614 |
| GSE220065 | Esophagus | Vercauteren<br>Drubbel and Beck. 2023<sup>63</sup> | 8 | GSM6784567,<br>GSM6784568,<br>GSM6784569 |
| GSE193376 | Esophagus | Kabir et al. 2022<sup>64</sup> | 12 | GSM5799214,<br>GSM5799218,<br>GSM5799222,<br>GSM5799226 |
| E-MTAB-8662 | Esophagus | McGinn et al. 2021<sup>65</sup> | 4,4,4,10,10,10 | P28_1, p28_2,<br>p28_3, Adult_1,<br>Adult_2, Adult_3 |
| GSE106273 | Mammary gland | Bach et al. 2017<sup>43</sup> | 8 | GSM2834498,<br>GSM2834499 |
| GSE150580 | Mammary gland | Li et al. 2020<sup>44</sup> | 12 | GSM4552664,<br>GSM4552665,<br>GSM4552666 |
| GSE164017 | Mammary gland | Cheng et al. 2023<sup>45</sup> | 2.5, 5, 5 | GSM4994963,<br>GSM4994964,<br>GSM4994965 |
| GSE191219 | Mammary gland | Saeki et al. 2024<sup>46</sup> | 43 | GSM5741807 |
| GSE120654 | Oral mucosa | Jones et al. 2019<sup>37</sup> | 12 | GSM3407039,<br>GSM3407040 |
| GSE228635 | Oral mucosa | Kondo et al. 2023<sup>38</sup> | 10 | GSM7134242 |
| GSE269575 | Oral mucosa | Jaber et al. 2024<sup>39</sup> | 8 | GSM8322397,<br>GSM8322398 |
| GSE146811 | Prostate | Karthaus et al. 2020<sup>66</sup> | 10 | GSM4474184,<br>GSM4474185,<br>GSM4474186,<br>GSM4474187,<br>GSM4474188,<br>GSM4474189,<br>GSM4474190 |
| GSE150692 | Prostate | Crowley et al. 2020<sup>67</sup> | 9 | GSM4556594,<br>GSM4556595,<br>GSM4556596, |
|  |  |  |  | GSM4556597,<br>GSM4556598,<br>GSM4556599 |
| GSE165741 | Prostate | Graham et al. 2023<sup>69</sup> | 24 | GSM5049201,<br>GSM5049204,<br>GSM5049207,<br>GSM5049210 |
| GSE224452 | Prostate | Aparicio et al. 2025<sup>68</sup> | 10 | GSM7024431,<br>GSM7024432 |
| GSE150327 | Salivary gland | Hauser et al. 2020<sup>51</sup> | 4, 4, 70 | GSM4546896,<br>GSM4546897,<br>GSM4546898 |
| GSE175649 | Salivary gland | Horeth et al. 2021<sup>52</sup> | 49 | GSM5343233 |
| GSE131339 | Thymus | Cowan et al. 2019<sup>40</sup> | 8 | GSM3770663,<br>GSM3770664,<br>GSM3770669,<br>GSM3770670 |
| GSE228198 | Thymus | Horie et al. 2023<sup>42</sup> | 9 | GSM7116509 |
| GSE240016 | Thymus | Kousa et al. 2024<sup>41</sup> | 8, 8, 8, 8, 72, 72,<br>72, 72 | GSM7679869,<br>GSM7679870,<br>GSM7679871,<br>GSM7679872,<br>GSM7679885,<br>GSM7679886,<br>GSM7679887,<br>GSM7679888 |
| GSE240020 | Thymus | Kousa et al. 2024<sup>41</sup> | 80 | GSM7963648 |
| GSE103354 | Trachea | Montoro et al. 2018<sup>57</sup> | 9 | GSM3314348,<br>GSM3314349,<br>GSM3314350,<br>GSM3314351,<br>GSM3314352,<br>GSM3314353 |
| GSE152692 | Trachea | Kiyokawa et al. 2021<sup>58</sup> | 8 | GSM4623795 |
| GSE244215 | Trachea | Lin et al. 2024<sup>59</sup> | 8, 96, 8, 96 | GSM7808936,<br>GSM7808937,<br>GSM7808938,<br>GSM7808939 |
| GSE145865 | Urethra | Joseph et al. 2020<sup>70</sup> | 10 | GSM4338168,<br>GSM4338169 |
| GSE173096 | Urethra | Joseph et al. 2021<sup>71</sup> | 10 | GSM5258273,<br>GSM5258274,<br>GSM5258275 |
| GSE201274 | Vagina | Kang et al. 2023<sup>47</sup> | 15 | GSM6057164 |
| E-MTAB-11491 | Vagina | Winkler et al. 2024<sup>48</sup> | 12, 12, 12, 12, 12,<br>12, 12, 12, 12, 12,<br>12, 12, 72, 72 | Ind001_vagina01,<br>Ind002_vagina02,<br>Ind003_vagina03,<br>Ind007_vagina07,<br>Ind007_vagina08,<br>Ind007_vagina09,<br>Ind007_vagina10,<br>Ind007_vagina11,<br>Ind007_vagina12,<br>Ind007_vagina13,<br>Ind007_vagina14,<br>Ind007_vagina15,<br>Ind001_vagina01,<br>Ind001_vagina01 |
| E-MTAB-11491 | Cervix | Winkler et al. 2024<sup>48</sup> | 12 | Ind001-cervix1,<br>Ind001-cervix2, |
|  |  |  |  | Ind001-cervix3,<br>Ind001-cervix4,<br>Ind001-cervix5,<br>Ind001-cervix6,<br>Ind001-cervix7,<br>Ind001-cervix8,<br>Ind001-cervix9,<br>Ind001-cervix10 |
| GSE128987 | Cervix | Chumduri et al. 2021<sup>49</sup> | 15 | GSM3689562 |
| GSE196529 | Cervix | Cooley et al. 2023<sup>50</sup> | 16 | GSM5888614 |
| GSE97213 | Epidermis, ChIP seq p63 | Fan et al. 2018<sup>74</sup> | New born | GSM2559737 |
|  | Epidermis, Drosopila melanogaster | Li et al. 2022<sup>89</sup> | Adult | body_wall_raw |
|  | Epidermis, Lamprey | Pang et al. 2025<sup>92</sup> | Larvae | Epidermis |
| GSE224695 | Epidermis, Danio rerio | Aman et al.<sup>94</sup> | 3 weeks PF | GSM7029635 |
| GSE195790 | Epidermis, Xenopus laevis | Liao et al. 2022<sup>93</sup> | Adult | GSM6214285 |
| GSE179690 | Epidermis, Gallus gallus | Lachner et al. 2022<sup>90</sup> | New born | GSM5428157 |
| GSE162185 | Epidermis, Homo sapiens | Gao et al. 2021<sup>91</sup> | Adult | GSM4946163 |

### Resources table

| REAGENT or RESOURCE | SOURCE | IDENTIFIER |
| --- | --- | --- |
| Antibodies |  |  |
| Rabbit anti-K5 | Covance | Cat# PRB-160P-100; RRID: AB_291581 |
| Chicken anti-K5 | Biologend | #905901 |
| Rat anti-K8 | Developmental Studies Hybridoma Bank, University of Iowa | Cat# TROMA-I; RRID: AB_531826 |
| Mouse anti-p63 | Abcam | Cat# ab735; RRID: AB_305870 |
| Chicken anti-K14 | Biologend | # 906004 |
| Rabbit anti-Ki67 | Invitrogen-Thermo Fisher Scientific | Cat# MA5-14520; RRID:AB_10979488 |
| Rabbit anti-HES1 | Cell Signaling technology | #11988 |
| Human anti-GFP | Institut Curie antibodies platform | NA |
| Rabbit anti-GFP | Institut Curie antibodies platform | NA |
| Guinea Pig anti-K13 | Progen | Cat# GP-K13; RRID: AB_1541010 |
| Guinea Pig anti-K10 | Progen | Cat# GP-K10; RRID: AB_2920688 |
| Rabbit anti-LORICRIN | Covance | Cat#: PRB-145P; RRID: AB_10064155 |
| Goat anti-rabbit AlexaFluor-coupled to different fluorochromes Cy5 | Invitrogen-Thermo Fisher Scientific | Cat#A10523; RRID: AB_2534032 |
| Goat anti-rabbit AlexaFluor-coupled to different fluorochromes A488 | Invitrogen-Thermo Fisher Scientific | Cat# A-11034; RRID: AB_2576217 |
| Goat anti-rabbit AlexaFluor-coupled to different fluorochromes A405 | Invitrogen-Thermo Fisher Scientific | Cat# A-31556; RRID: AB_221605 |
| Goat anti-mouse AlexaFluor-coupled to different fluorochromes Cy3 | Invitrogen-Thermo Fisher Scientific | Cat# A10521; RRID: AB_2534030 |
| Goat anti-mouse AlexaFluor-coupled to different fluorochromes Cy5 | Invitrogen-Thermo Fisher Scientific | Cat# A10524; RRID: AB_2534033 |
| Goat anti-mouse AlexaFluor-coupled to different fluorochromes A488 | Invitrogen-Thermo Fisher Scientific | Cat# A-11001; RRID: AB_2534069 |
| Goat anti-rat AlexaFluor-coupled to different fluorochromes Cy3 | Invitrogen-Thermo Fisher Scientific | Cat# A10522; RRID: AB_1500704 |
| Goat anti-rat AlexaFluor-coupled to different fluorochromes Cy5 | Invitrogen-Thermo Fisher Scientific | Cat# A1052; RRID: AB_10562712 |
| Goat anti-rat AlexaFluor-coupled to different fluorochromes A488 | Invitrogen-Thermo Fisher Scientific | Cat# A-11006; RRID: AB_141373 |
| Goat anti-guinea pig AlexaFluor-coupled to different fluorochromes (A568) | Invitrogen-Thermo Fisher Scientific | CAT#A-11075; RRID: AB_2337423 |
| Chemicals, peptides, and recombinant proteins |  |  |
| Sunflower Oil | Merck-Millipore | S5007 |
| Tamoxifen | MP Biomedicals | 0215673891 |
| Paraformaldehyde (PFA) | Electron Microscopy Sciences | 15710 |
| Aqua poly/mount | Tebu Bio | 07918606-5 |
| Glycerol | Euromedex | 15710 |
| Triton X-100 | Euromedex | 2000C |
| Sucrose | Sigma | S0389 |
| Tissue-Tek O.C.T. | Sakura | 4583 |
| Experimental models: Organisms/strains |  |  |
| *K5-CreERT2* | Van Keymeulen et al.<sup>8</sup> | RRID:IMSR_JAX:029155, |
| *R26-N1ICD* | Murtaugh et al.<sup>81</sup> | RRID:IMSR_JAX:008159 |
| *R26-mTmG* | Muzumdar et al.<sup>71</sup> | RRID:IMSR_JAX:037456 |
| Software and algorithms |  |  |
| R v4.0.3 | R Core Team | https://www.r-project.org/ |
| Python v3.7.7 | Python Software Foundation | https://python.org/ |
| Seurat v4.2.0 | Satija Lab | https://satijalab.org/seurat/ |
| ImageJ | National Institutes of Health | https://imagej.nih.gov/ij/ |
| Zen blue 3.5 (ZEISS ZEN lite) | Carl Zeiss Microscopy | https://www.zeiss.com/microscopy/en/products/software/zeiss-zen-lite.html |

