## Supplementary material for "Single-cell transcriptomics uncover conserved molecular mechanisms and functional diversification in multilayered epithelia": Supplemntary figures

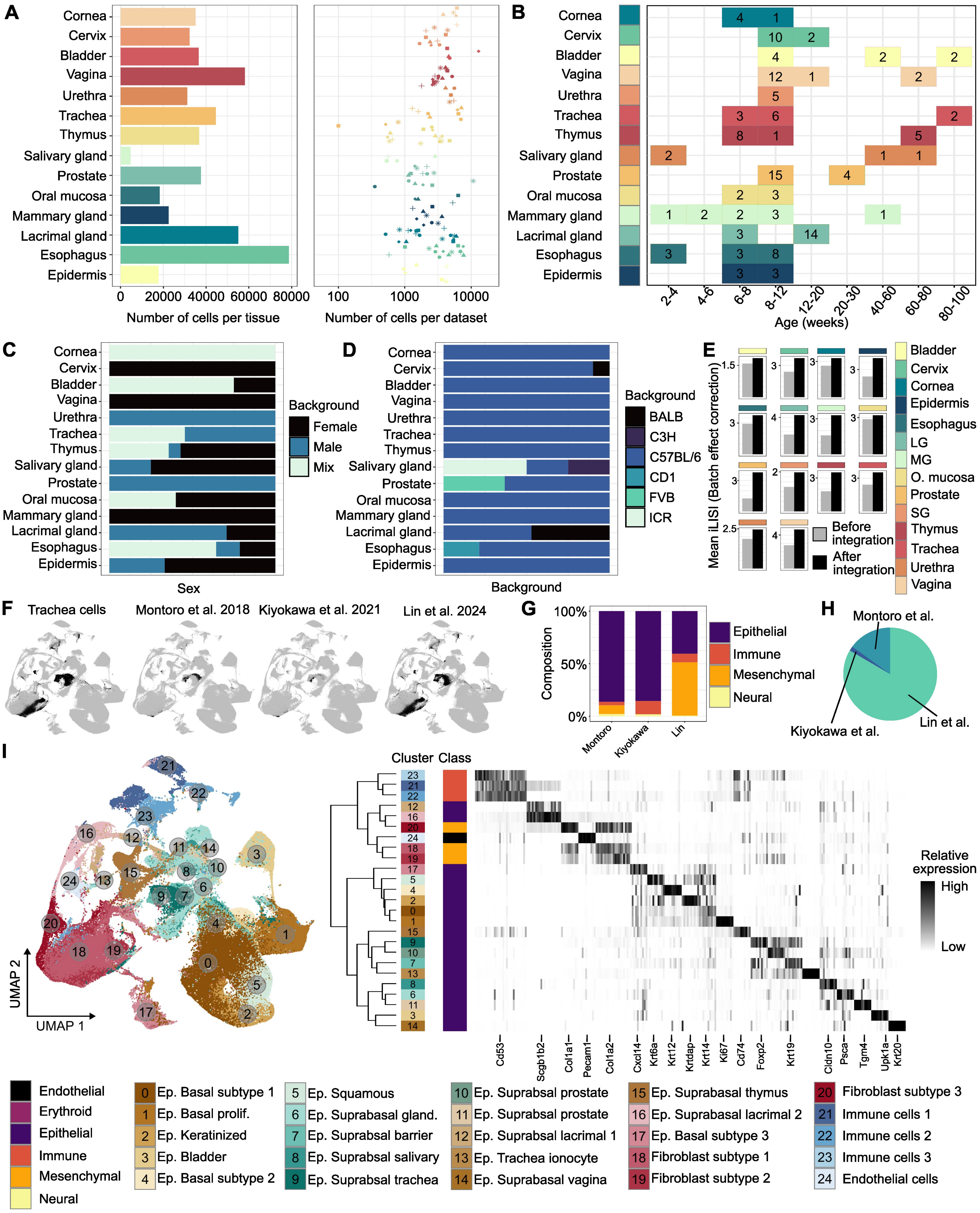

A
Cornea
Cervix
Bladder
Vagina
Urethra
Trachea
Thymus
Salivary gland
Prostate
Oral mucosa
Mammary gland
Lacrimal gland
Esophagus
Epidermis
0 20000 40000 60000 80000
Number of cells per tissue
100 1000 10000
Number of cells per dataset
B
2-4 4-6 6-8 8-12 12-20 20-30 40-60 60-80 80-100
Age (weeks)
C
Background
Female
Male
Mix
Sex
D
Background
BALB
C3H
C57BL/6
CD1
FVB
ICR
E
Mean iLISI (Batch effect correction)
Before integration
After integration
Bladder
Cervix
Cornea
Epidermis
Esophagus
LG
MG
O. mucosa
Prostate
SG
Thymus
Trachea
Urethra
Vagina
F
Trachea cells
Montoro et al. 2018
Kiyokawa et al. 2021
Lin et al. 2024
G
Composition
Epithelial
Immune
Mesenchymal
Neural
Montoro
Kiyokawa
Lin
H
Montoro et al.
Kiyokawa et al.
Lin et al.
I
UMAP 1
UMAP 2
Cluster
Class
Relative expression
High
Low
Cd53
Scgb1b2
Col1a1
Pecam1
Col1a2
Cxcl14
Krt6a
Krt12
Krtdap
Krt14
Ki67
Cd74
Foxp2
Krt19
Cldn10
Psca
Tgm4
Upk1a
Krt20
Endothelial
Erythroid
Epithelial
Immune
Mesenchymal
Neural
0 Ep. Basal subtype 1
1 Ep. Basal prolif.
2 Ep. Keratinized
3 Ep. Bladder
4 Ep. Basal subtype 2
5 Ep. Squamous
6 Ep. Suprabasal gland.
7 Ep. Suprabsal barrier
8 Ep. Suprabsal salivary
9 Ep. Suprabsal trachea
10 Ep. Suprabsal prostate
11 Ep. Suprabsal prostate
12 Ep. Suprabsal lacrimal 1
13 Ep. Trachea ionocyte
14 Ep. Suprabasal vagina
15 Ep. Suprabasal thymus
16 Ep. Suprabasal lacrimal 2
17 Ep. Basal subtype 3
18 Fibroblast subtype 1
19 Fibroblast subtype 2
20 Fibroblast subtype 3
21 Immune cells 1
22 Immune cells 2
23 Immune cells 3
24 Endothelial cells

***Figure S1. Atlas. Related to Figure 1.***

*A: Number of cells, after quality filtering, per tissue (left) and per dataset (right). B: Age of mice (in weeks) associated with the individual datasets included in the atlas. C: Proportion of cells coming from scRNA seq dataset composed of male, female or a mix of both sex for each tissue. D: Proportion of cells from each mouse genetic background for each tissue. E: Evaluation of batch effect correction through LISI before and after CSS integration (See Methods). F: Projection of tracheal cells onto the integrated epithelial atlas. Cells from different studies are treated as independent biological replicates. G: Composition of the three tracheal scRNA-seq datasets included in the analysis. H: Relative contribution of each dataset to the total number of tracheal cells. I: Integrated UMAP of scRNA-seq data colored by cell clusters and associated heatmap displaying the top 10 cluster specific markers.*

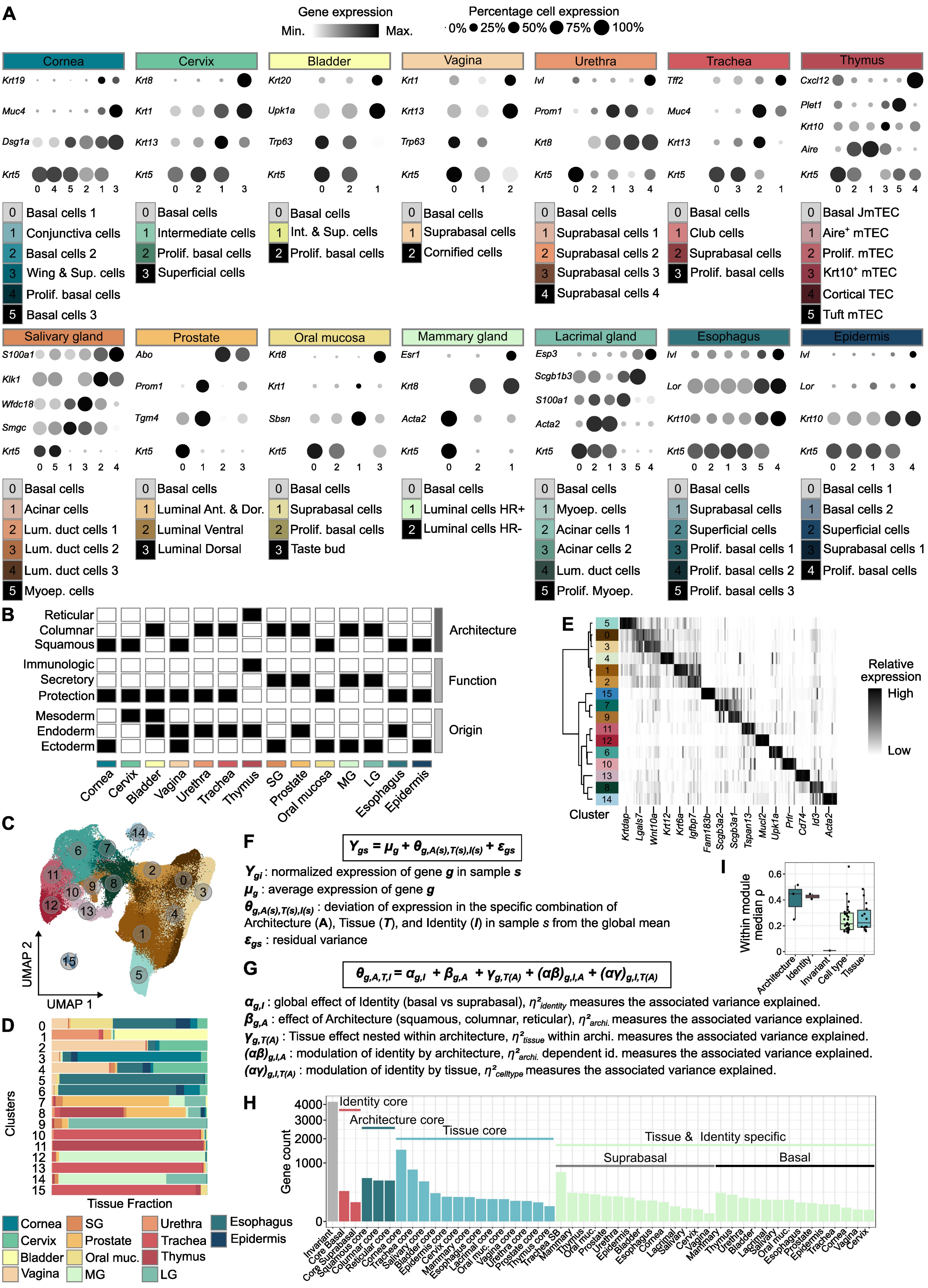
A
Gene expression
Min.
Max.
Percentage cell expression
0% 25% 50% 75% 100%
Cornea
Cervix
Bladder
Vagina
Urethra
Trachea
Thymus
Basal cells 1
Conjunctiva cells
Basal cells 2
Wing & Sup. cells
Prolif. basal cells
Basal cells 3
Basal cells
Intermediate cells
Prolif. basal cells
Superficial cells
Int. & Sup. cells
Suprabasal cells
Cornified cells
Suprabasal cells 1
Suprabasal cells 2
Suprabasal cells 3
Suprabasal cells 4
Club cells
Basal JmTEC
Aire+ mTEC
Prolif. mTEC
Krt10+ mTEC
Cortical TEC
Tuft mTEC
Salivary gland
Prostate
Oral mucosa
Mammary gland
Lacrimal gland
Esophagus
Epidermis
Acinar cells
Lum. duct cells 1
Lum. duct cells 2
Lum. duct cells 3
Myoep. cells
Luminal Ant. & Dor.
Luminal Ventral
Luminal Dorsal
Taste bud
Luminal cells HR+
Luminal cells HR-
Acinar cells 1
Acinar cells 2
Lum. duct cells
Prolif. Myoep.
Prolif. basal cells 1
Prolif. basal cells 2
Prolif. basal cells 3
Superficial cells
B
Reticular
Columnar
Squamous
Architecture
Immunologic
Secretory
Protection
Function
Mesoderm
Endoderm
Ectoderm
Origin
Cornea Cervix Bladder Vagina Urethra Trachea Thymus SG Prostate Oral mucosa MG LG Esophagus Epidermis
C
UMAP 1
UMAP 2
D
Clusters
Tissue Fraction
Cornea SG Urethra Esophagus
Cervix Prostate Trachea Epidermis
Bladder Oral muc. Thymus
Vagina MG LG
E
Relative expression
High
Low
Cluster
Krtdap Lgals7 Wnt10a Krt12 Krt6a Igfbp7 Fam183b Scgb3a2 Scgb3a1 Tspan13 Mucl2 Upk1a Prl Cd74 Id3 Acta2
F
$Y_{gs} = \mu_g + \theta_{g,A(s),T(s),I(s)} + \varepsilon_{gs}$
$Y_{gi}$ : normalized expression of gene $g$ in sample $s$
$\mu_g$ : average expression of gene $g$
$\theta_{g,A(s),T(s),I(s)}$ : deviation of expression in the specific combination of Architecture (A), Tissue (T), and Identity (I) in sample s from the global mean
$\varepsilon_{gs}$ : residual variance
G
$\theta_{g,A,T,I} = \alpha_{g,I} + \beta_{g,A} + \gamma_{g,T(A)} + (\alpha\beta)_{g,I,A} + (\alpha\gamma)_{g,I,T(A)}$
$\alpha_{g,I}$ : global effect of Identity (basal vs suprabasal), $\eta^2_{identity}$ measures the associated variance explained.
$\beta_{g,A}$ : effect of Architecture (squamous, columnar, reticular), $\eta^2_{archi.}$ measures the associated variance explained.
$\gamma_{g,T(A)}$ : Tissue effect nested within architecture, $\eta^2_{tissue}$ within archi. measures the associated variance explained.
$(\alpha\beta)_{g,I,A}$ : modulation of identity by architecture, $\eta^2_{archi.}$ dependent id. measures the associated variance explained.
$(\alpha\gamma)_{g,I,T(A)}$ : modulation of identity by tissue, $\eta^2_{celltype}$ measures the associated variance explained.
H
Gene count
Identity core
Architecture core
Tissue core
Tissue & Identity specific
Suprabasal
Basal
I
Within module median ρ
Architecture Identity Invariant Cell type Tissue

***Figure S2. Atlas. Related to Figure 2.***

***A:*** *Dot plot showing the average gene expression of established cell type markers across epithelial cell populations.*

***B:*** *Table recapitulating the Origin, Physiological Function and Tissue Architecture of each of the 14 multilayered murine epithelia.* ***C-E:*** *Integrated UMAP of epithelial scRNA-seq data colored by cell clusters (B), tissue fraction for each tissue within each cluster of the epithelial integrated atlas (C) and heatmap displaying the top 10 cluster specific markers (D).* ***F-G****: Variance decomposition model. For each gene g, expression across samples s was modeled as the sum of a global mean ($\mu_g$) and a deviation term $\theta_{g,A(s),T(s),I(s)}$ reflecting the combined effects of epithelial Architecture (A), Tissue (T), and Identity (I), plus residual variance $\varepsilon_{gs}$. Each deviation term was further decomposed into additive and interaction components: a global identity effect ($\alpha_{g,I}$), an architectural effect ($\beta_{g,A}$), a tissue effect nested within architecture ($\gamma_{g,T(A)}$), and interaction terms describing modulation of identity by architecture or tissue $(\alpha\beta)_{g,I,A}$ and $(\alpha\gamma)_{g,I,T(A)}$. This hierarchical linear model quantifies how much of each gene's expression variance is explained by identity-, architecture-, and tissue-level factors, revealing the nested structure of epithelial transcriptomic organization (See Methods).* ***H****: Bar plot showing number of genes assigned to each module following the decision tree presented in the method section.* ***I:*** *Intra-module average correlation coefficient for each category of module.*

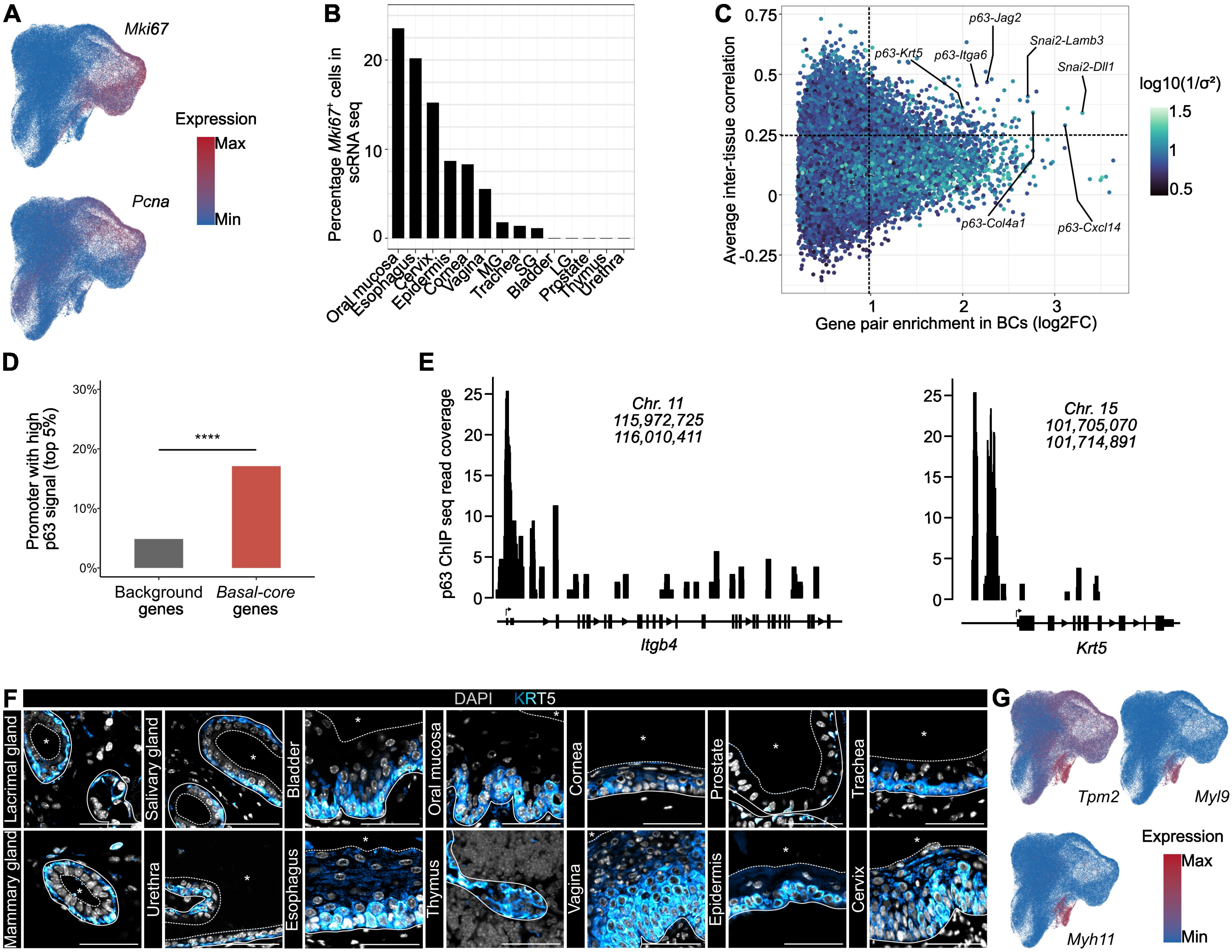

A
Mki67
Pcna
Expression
Max
Min
B
Percentage Mki67+ cells in scRNA seq
Oral mucosa
Esophagus
Cervix
Epidermis
Cornea
Vagina
MG
Trachea
SG
Bladder
LG
Prostate
Thymus
Urethra
C
Average inter-tissue correlation
Gene pair enrichment in BCs (log2FC)
p63-Jag2
Snai2-Lamb3
p63-Krt5
p63-Itga6
Snai2-Dll1
p63-Col4a1
p63-Cxcl14
log10(1/σ²)
D
Promoter with high p63 signal (top 5%)
****
Background genes
Basal-core genes
E
p63 ChIP seq read coverage
Chr. 11
115,972,725
116,010,411
Itgb4
Chr. 15
101,705,070
101,714,891
Krt5
F
DAPI KRT5
Lacrimal gland
Salivary gland
Bladder
Oral mucosa
Cornea
Prostate
Trachea
Mammary gland
Urethra
Esophagus
Thymus
Vagina
Epidermis
Cervix
G
Tpm2
Myl9
Myh11
Expression
Max
Min

***Figure S3. Atlas. Related to Figure 3.***

***A:*** *Integrated UMAP of BCs scRNA-seq data colored by expression level of Mki67 and Pcna.* ***B:*** *Percentage of Mki67+ cells per tissue in the scRNA seq atlas.* ***C:*** *Scatter plot showing co-expressed transcription factor–containing gene pairs that are consistently positively correlated across multiple stratified epithelial tissues. Each point represents a gene pair, plotted by its mean* $log_2$ *fold-change between basal and suprabasal compartments (x-axis) and average co-expression strength across tissues (y-axis). Points are colored by the* $log_{10}$*-transformed precision of the co-expression estimate (inverse of variance across tissues).* ***D****: Bar plot showing the fraction of promoters with high p63 ChIP-seq signal (≥95th percentile of genome-wide signal) in basal-core genes (red) versus background promoters (gray).* ***E:*** *ΔNp63 ChIP-seq read coverage across Itgb4 and Krt5 loci (mm10 assembly). Signal tracks represent aligned ChIP-seq raw coverage for ΔNp63 in mouse keratinocytes.* ***F:*** *Confocal imaging of adult WT multilayered epithelia stained for Dapi and KRT5 (Cyan).* ***G:*** *Integrated UMAP of BCs scRNA-seq data colored by expression level of genes associated with contractility Tpm2, Myl9 and Myh11. *: indicate lumen or external space; dotted line: indicate the interface between apical domain of epithelia and external space; continuous line: indicate the interface between epithelium and basement membrane. Scale bar: 50 µm.*

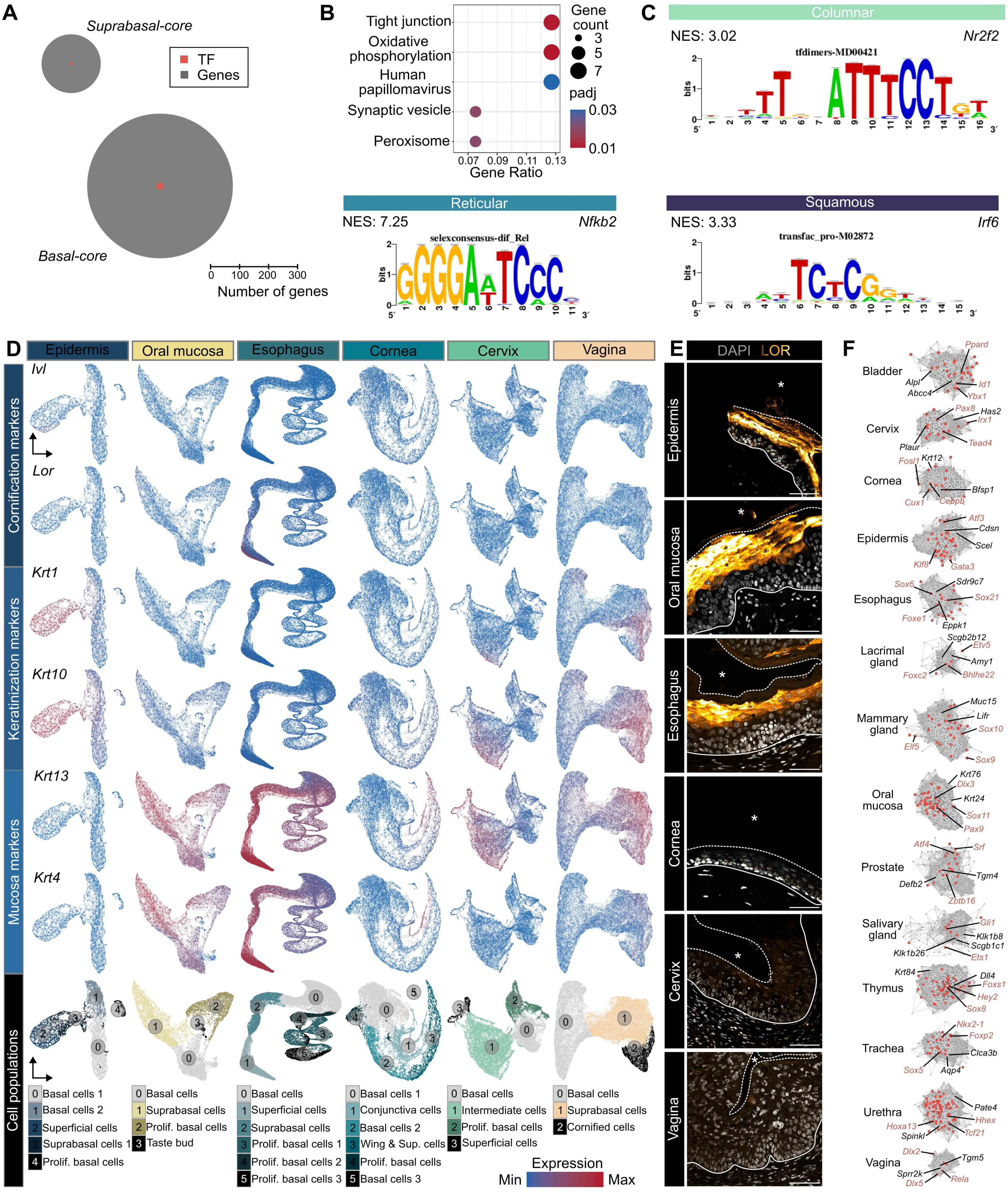

A
Suprabasal-core
TF
Genes
Basal-core
0 100 200 300
Number of genes
B
Tight junction
Oxidative phosphorylation
Human papillomavirus
Synaptic vesicle
Peroxisome
0.07 0.09 0.11 0.13
Gene Ratio
Gene count
3
5
7
padj
0.03
0.01
C
Columnar
NES: 3.02
Nr2f2
tfdimers-MD00421
Reticular
NES: 7.25
Nfkb2
selexconsensus-dif_Rel
Squamous
NES: 3.33
Irf6
transfac_pro-M02872
bits
D
Epidermis
Oral mucosa
Esophagus
Cornea
Cervix
Vagina
Cornification markers
Ivl
Lor
Keratinization markers
Krt1
Krt10
Mucosa markers
Krt13
Krt4
Cell populations
0 Basal cells 1
1 Basal cells 2
2 Superficial cells
3 Suprabasal cells 1
4 Prolif. basal cells
0 Basal cells
1 Suprabasal cells
2 Prolif. basal cells
3 Taste bud
0 Basal cells
1 Superficial cells
2 Suprabasal cells
3 Prolif. basal cells 1
4 Prolif. basal cells 2
5 Prolif. basal cells 3
0 Basal cells 1
1 Conjunctiva cells
2 Basal cells 2
3 Wing & Sup. cells
4 Prolif. basal cells
5 Basal cells 3
0 Basal cells
1 Intermediate cells
2 Prolif. basal cells
3 Superficial cells
0 Basal cells
1 Suprabasal cells
2 Cornified cells
Expression
Min
Max
E
DAPI LOR
Epidermis
Oral mucosa
Esophagus
Cornea
Cervix
Vagina
F
Bladder
Ppard
Alpl
Abcc4
Id1
Ybx1
Cervix
Pax8
Has2
Irx1
Plaur
Tead4
Cornea
Fosl1
Krt12
Bfsp1
Cux1
Cebpb
Epidermis
Atf3
Cdsn
Scel
Klf8
Gata3
Esophagus
Sox6
Sdr9c7
Sox21
Foxe1
Eppk1
Lacrimal gland
Scgb2b12
Etv5
Amy1
Foxc2
Bhlhe22
Mammary gland
Muc15
Lifr
Sox10
Elf5
Sox9
Oral mucosa
Krt76
Dlx3
Krt24
Sox11
Pax9
Prostate
Atf4
Srf
Tgm4
Defb2
Zbtb16
Salivary gland
Gli1
Klk1b8
Scgb1c1
Klk1b26
Ets1
Thymus
Krt84
Dll4
Foxs1
Hey2
Sox8
Trachea
Nkx2-1
Foxp2
Clca3b
Sox5
Aqp4
Urethra
Pate4
Hhex
Hoxa13
Tcf21
Spinkl
Vagina
Dlx2
Tgm5
Sprr2k
Dlx5
Rela

***Figure S4. Atlas. Related to Figure 4.***

***A:*** *Number of genes contained in Basal core and Suprabasal core modules.* ***B:*** *KEGG enriched terms in the Suprabasal core module.* ***C****: Motif enrichment of architecture-specific transcription factors. Representative transcription factor motifs identified by RcisTarget analysis in genes belonging to architecture-specific modules. The plots show the top-scoring motifs enriched within ±1 kb of transcription start sites for Columnar (Nr2f2), Reticular (Nfkb2) and Squamous (Irf6) modules, with normalized enrichment scores (NES) indicated. Each motif represents the consensus sequence most strongly enriched in the promoters of genes regulated within the corresponding architectural program (See Methods).* ***D:*** *UMAP of individual squamous epithelia colored by expression of genes associated with cornification, keratinization or mucosal tissues.* ***E:*** *Confocal imaging of cross section of adult WT murine tissues stained for Dapi and LORICRIN (LOR). *: indicate lumen or external space.* ***F:*** *Tissue core GRN modules and associated regulators. Scale bar: 50 µm.*

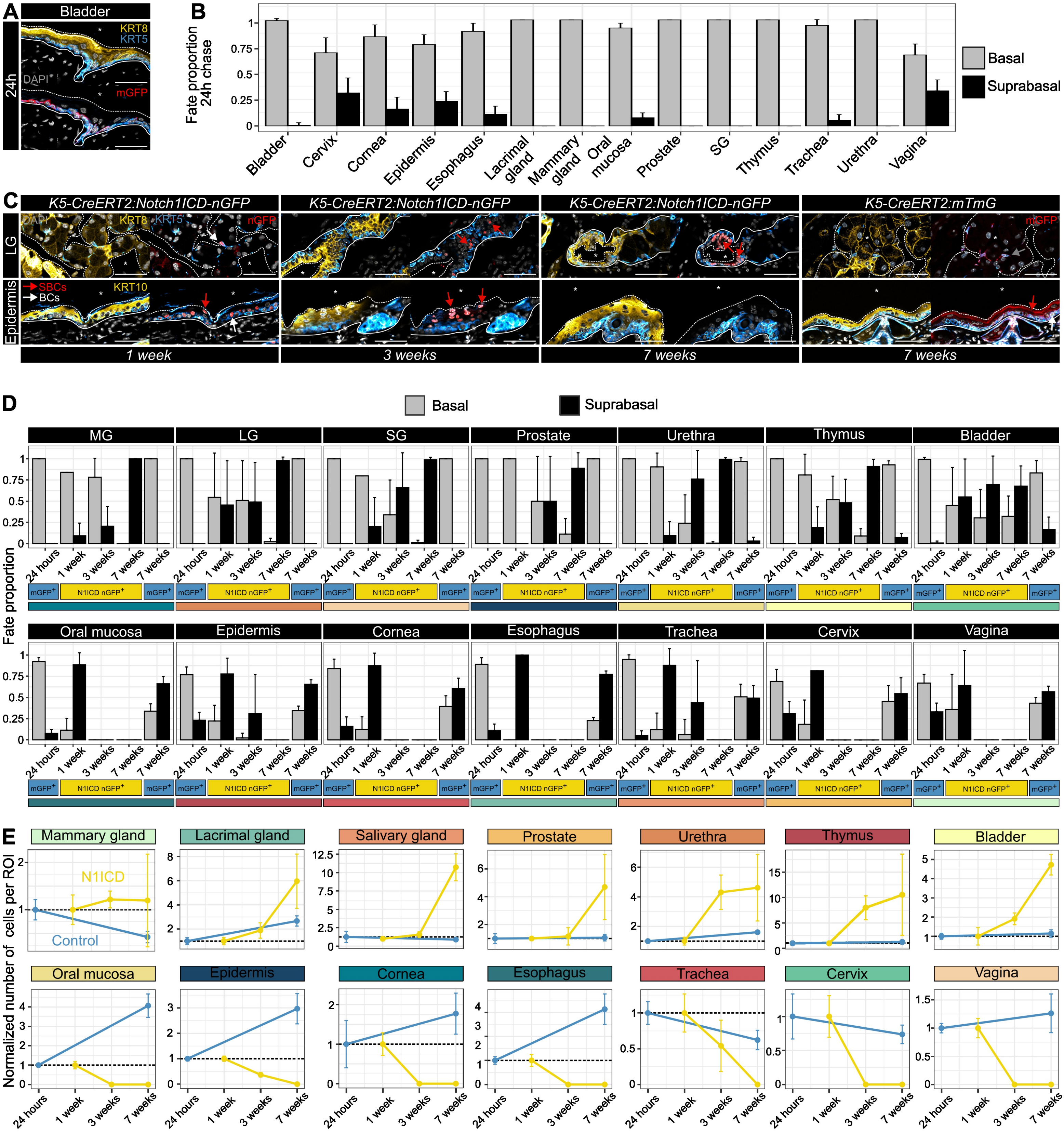

A
Bladder
24h
KRT8
KRT5
DAPI
mGFP
B
Fate proportion 24h chase
Basal
Suprabasal
Bladder
Cervix
Cornea
Epidermis
Esophagus
Lacrimal gland
Mammary gland
Oral mucosa
Prostate
SG
Thymus
Trachea
Urethra
Vagina
C
K5-CreERT2:Notch1ICD-nGFP
K5-CreERT2:mTmG
LG
Epidermis
nGFP
KRT10
SBCs
BCs
1 week
3 weeks
7 weeks
D
Fate proportion
MG
LG
SG
Prostate
Urethra
Thymus
Bladder
Oral mucosa
Epidermis
Cornea
Esophagus
Trachea
Cervix
Vagina
24 hours
1 week
3 weeks
7 weeks
mGFP+
N1ICD nGFP+
E
Normalized number of cells per ROI
Mammary gland
Lacrimal gland
Salivary gland
Prostate
Urethra
Thymus
Bladder
Oral mucosa
Epidermis
Cornea
Esophagus
Trachea
Cervix
Vagina
N1ICD
Control

***Figure S5. Atlas. Related to Figure 5.***

***A:*** *Confocal imaging of immunostaining for K5, K8 and mGFP Bladder from K5-Cre$^{ERT2}$:mTmG mouse after 24h of lineage tracing.* ***B:*** *Quantification of fate of mGFP$^{+}$ cells after 24 hours of lineage tracing. Data are displayed as Mean + s.d. Chi-squared test (n=3 biological replicates).* ***C:*** *Confocal imaging of immunostaining for KRT5, suprabasal marker (KRT8 or KRT10) and mGFP in lacrimal gland (LG) and Epidermis from K5-Cre$^{ERT2}$:mTmG or K5-Cre$^{ERT2}$:N1ICD-nls-GFP mouse after 1, 3 or 7 weeks of lineage tracing.* ***D:*** *Quantification of fate of mGFP$^{+}$ or nGFP$^{+}$ cells after different chase of lineage tracing. Data are displayed as Mean + s.d. Chi-squared test (n=3 biological replicates).* ***E:*** *Normalized GFP$^{+}$ cells per ROI after different chase. Data are normalized to the mean number of GFP+ cells observed at the earlier chase timepoint (24h mGFP$^{+}$, 1-week nGFP$^{+}$) Scale bar: 50 µm.*

A

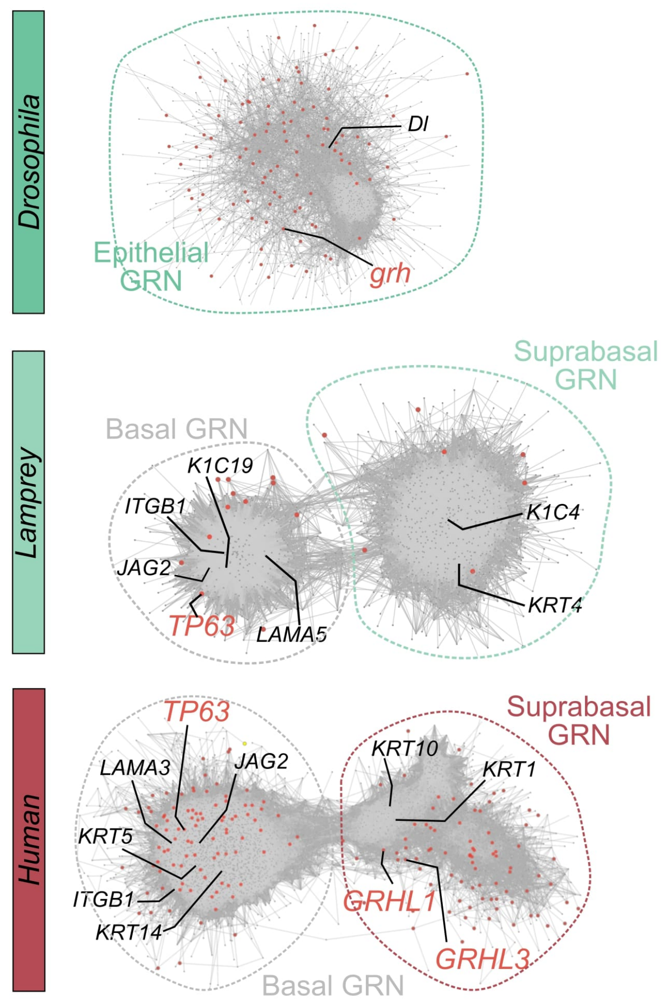


B

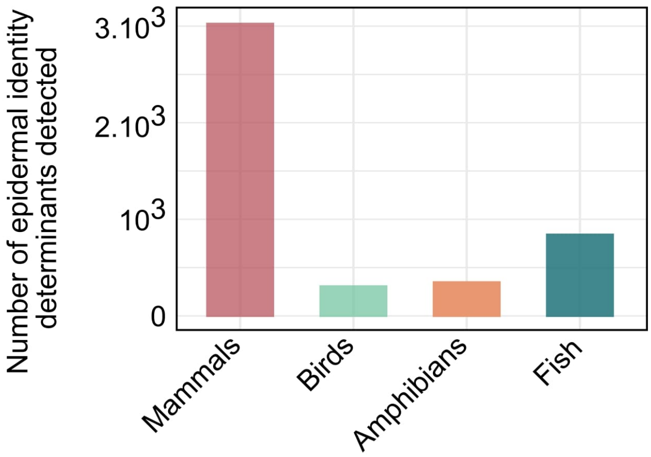


C

| Shared | Mammals | Fish | Birds | Amphibians |
|---|---|---|---|---|
| *Krt5* | *Sprr1b* | *muc5a* | *EDKMB* | *tuba1c.L* |
| *Col17a1* | *Sprr2a* | *muc13a* | *EDQ1* | *tuba3c.S* |
| *Itgb1* | *Sprr2d* | *muc2.2* | *EDQM1* | Ciliated cells marker |
| *Sparc* | *Sprrd1* | *muc5.2* | *EDQM2* | |
| *Jag2* | *Sprr2e* | Mucus secreted proteins | Bird specific *stratum corneum* compounds | |
| *Fgfr2* | Mammal specific *stratum corneum* proteins | | | |
| *Fermt1* | | | | |
| *Macf1* | | | | |
| *Rab10* | | | | |
| *Atp1a1* | | | | |
| *Trp63* | | | | |
| *Tpt1* | | | | |
| *Net1* | | | | |
| *Fam212a* | | | | |
| *Hspa8* | | | | |
| *Tpm3* | | | | |
| *Cnpb* | | | | |

***Figure S6. Atlas. Related to Figure 6.***

***A:*** *Gene co-expression network in epidermis scRNA seq dataset of Drosophila, Lamprey and Human.* ***B:*** *Number of genes composing basal and suprabasal GRN for each species.* ***C:*** *Example of genes shared across species or species specific.*

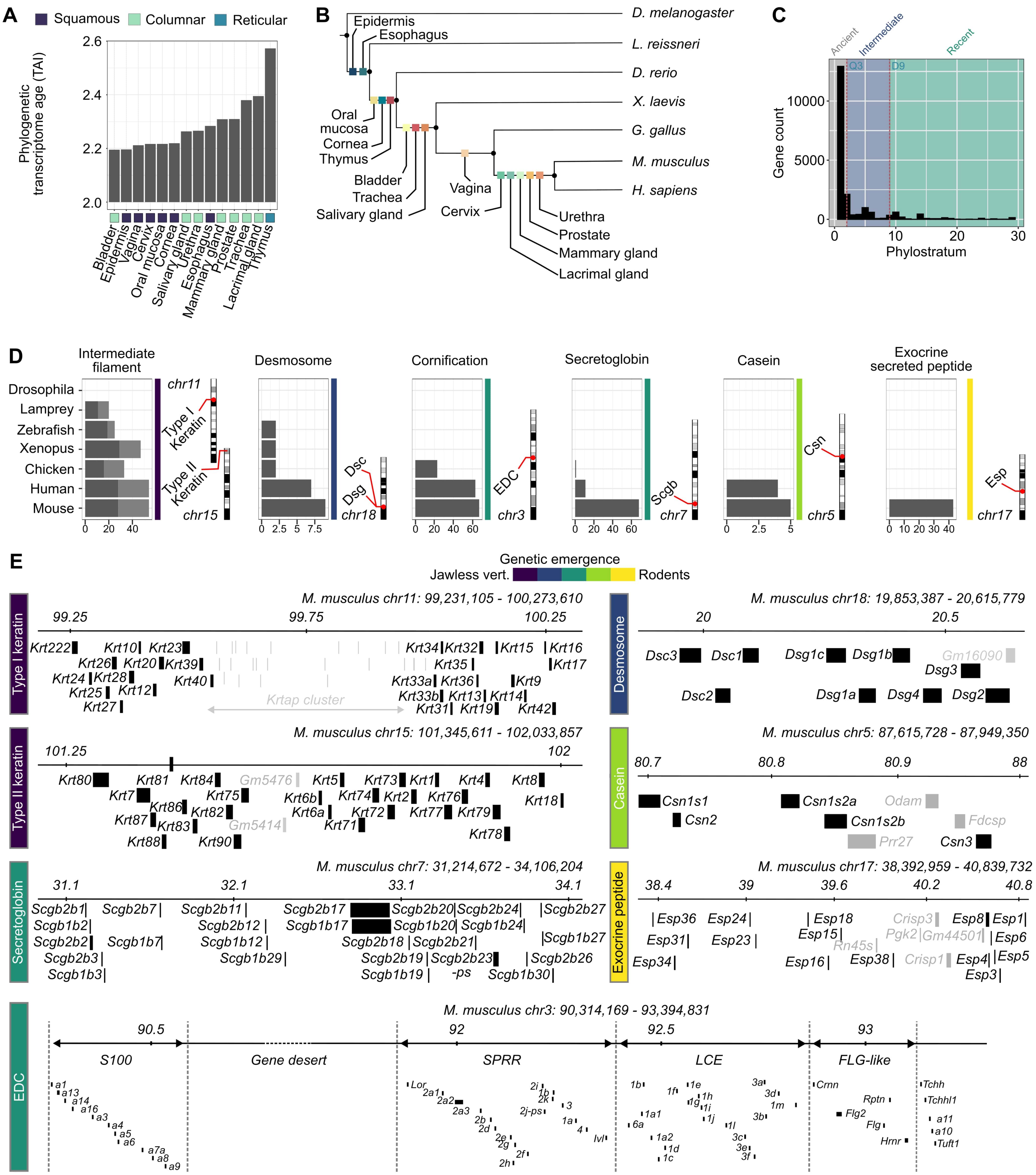

A
Squamous
Columnar
Reticular
Phylogenetic transcriptome age (TAI)
2.6
2.4
2.2
2.0
Bladder
Epidermis
Vagina
Cervix
Oral mucosa
Cornea
Salivary gland
Urethra
Esophagus
Mammary gland
Prostate
Trachea
Lacrimal gland
Thymus
B
Epidermis
Esophagus
Oral mucosa
Cornea
Thymus
Bladder
Trachea
Salivary gland
Vagina
Cervix
Urethra
Prostate
Mammary gland
Lacrimal gland
D. melanogaster
L. reissneri
D. rerio
X. laevis
G. gallus
M. musculus
H. sapiens
C
Ancient
Intermediate
Recent
Q3
D9
Gene count
10000
5000
0
Phylostratum
0
10
20
30
D
Intermediate filament
Desmosome
Cornification
Secretoglobin
Casein
Exocrine secreted peptide
Drosophila
Lamprey
Zebrafish
Xenopus
Chicken
Human
Mouse
chr11
Type I Keratin
Type II Keratin
chr15
Dsc
Dsg
chr18
EDC
chr3
Scgb
chr7
Csn
chr5
Esp
chr17
E
Genetic emergence
Jawless vert.
Rodents
Type I keratin
M. musculus chr11: 99,231,105 - 100,273,610
99.25
99.75
100.25
Krt222 Krt10 Krt23 Krt34 Krt32 Krt15 Krt16
Krt26 Krt20 Krt39 Krt35 Krt17
Krt24 Krt28 Krt40 Krt33a Krt36 Krt9
Krt25 Krt12 Krt33b Krt13 Krt14
Krt27 Krt31 Krt19 Krt42
Krtap cluster
Desmosome
M. musculus chr18: 19,853,387 - 20,615,779
20
20.5
Dsc3 Dsc1 Dsg1c Dsg1b Gm16090
Dsg3
Dsc2 Dsg1a Dsg4 Dsg2
Type II keratin
M. musculus chr15: 101,345,611 - 102,033,857
101.25
102
Krt80 Krt81 Krt84 Gm5476 Krt5 Krt73 Krt1 Krt4 Krt8
Krt7 Krt75 Krt6b Krt74 Krt2 Krt76 Krt18
Krt86 Krt82 Krt6a Krt72 Krt77 Krt79
Krt87 Krt83 Gm5414 Krt71
Krt88 Krt90 Krt78
Casein
M. musculus chr5: 87,615,728 - 87,949,350
80.7
80.8
80.9
88
Csn1s1 Csn1s2a Odam
Csn2 Csn1s2b Fdcsp
Prr27 Csn3
Secretoglobin
M. musculus chr7: 31,214,672 - 34,106,204
31.1
32.1
33.1
34.1
Scgb2b1 Scgb2b7 Scgb2b11 Scgb2b17 Scgb2b20 Scgb2b24 Scgb2b27
Scgb1b2 Scgb2b12 Scgb1b17 Scgb1b20 Scgb1b24
Scgb2b2 Scgb1b7 Scgb1b12 Scgb2b18 Scgb2b21 Scgb1b27
Scgb2b3 Scgb1b29 Scgb2b19 Scgb2b23 Scgb2b26
Scgb1b3 Scgb1b19 -ps Scgb1b30
Exocrine peptide
M. musculus chr17: 38,392,959 - 40,839,732
38.4
39
39.6
40.2
40.8
Esp36 Esp24 Esp18 Crisp3 Esp8 Esp1
Esp15 Pgk2 Gm44501 Esp6
Esp31 Esp23 Rn45s
Esp34 Esp16 Esp38 Crisp1 Esp4 Esp5
Esp3
EDC
M. musculus chr3: 90,314,169 - 93,394,831
90.5
92
92.5
93
S100
Gene desert
SPRR
LCE
FLG-like
a1 a13 a14 a16 a3 a4 a5 a6 a7a a8 a9
Lor 2a1 2a2 2a3 2b 2d 2e 2g 2h 2f 2i 1b 2k 2j-ps 3 1a 4 Ivl
1b 1f 1e 1g 1h 1i 1j 1a1 6a 1a2 1d 1c 1l 3a 3d 1m 3b 3c 3e 3f
Crnn Rptn Flg2 Flg Hrnr
Tchh Tchhl1 a11 a10 Tuft1

***Figure S7. Atlas. Related to Figure 7.***

***A:*** *Phylogenetic transcriptome age index (TAI) for each murine tissue.* ***B:*** *Phylogenetic tree of animal models studied and associated multilayered epithelia emergence.* ***C:*** *Distribution of mouse genome wide murine phylostrata. 3$^{rd}$ quartile and (Q3) and 9$^{th}$ decile (D9) are marked by vertical lines. Classification in Ancient, Intermediate or Recent is based on these thresholds.* ***D:*** *Genomic localization on the mm10 genome and number of genes part of each gene cluster across organisms.* ***E:*** *Organization of gene clusters within the Mus musculus (mm10) genome assembly.*
